# Unraveling an unknown diversity of archaeal and bacterial tetraether membrane lipid producers in a euxinic marine system

**DOI:** 10.1101/2024.06.25.600576

**Authors:** Dina Castillo Boukhchtaber, F. A. Bastiaan von Meijenfeldt, Diana X. Sahonero Canavesi, Denise Dorhout, Nicole J. Bale, Ellen C. Hopmans, Laura Villanueva

## Abstract

Bacterial membrane lipids have been traditionally defined as fatty acids (FAs) bilayers linked through ester bonds, while those of Archaea as ether-linked isoprenoids forming bilayers or monolayers of membrane spanning lipids (MSLs) known as isoprenoidal glycerol dialkyl glycerol tetraethers (isoGDGTs). This paradigm has been challenged with the discovery of branched GDGTs (brGDGTs), membrane spanning ether-bound branched alkyl FAs, that are of bacterial origin but whose specific producers in the environment are often unknown. The limited number of available microbial cultures restricts the knowledge of the biological sources of membrane lipids, which in turn limits their potential applicability as biomarkers. To address this limitation, we detected membrane lipids in the Black Sea using high resolution accurate mass/mass spectrometry and inferred their potential producers by targeting lipid biosynthetic pathways encoded on the metagenome, in metagenome-assembled genomes and unbinned scaffolds. We also detected brGDGTs and overly branched GDGTs in the suboxic and euxinic waters, which are potentially attributed, to members of the Planctomycetota, Cloacimonadota, Desulfobacterota, Chloroflexota, Actinobacteria and Myxococcota—all anaerobic microorganisms. These results open a new chapter in the use of specific brGDGTs as biomarkers of anoxic conditions in marine settings and of the role of these membrane lipids in microbial adaptation.

## Introduction

Microbial membrane lipids are the main components of the cell membrane, which are either composed of fatty acids linked through ester bonds to glycerol-3-phosphate (G3P) in the case of bacteria, or of isoprenoids linked through ether bonds to glycerol-1-phosphate (G1P) in archaea. Archaeal membranes can be organized as bilayers of diether lipids or as monolayers of tetraethers or so-called isoprenoid glycerol dialkyl glycerol tetraethers (isoGDGTs) (Koga and Morii, 2007; Villanueva *et al*., 2014). Conversely, bacterial membranes are generally composed of bilayers. Exceptions are bacterial membrane spanning lipids (MSLs) composed of branched alkyl chains derived from fatty acids (FAs) and ether bonds, or so-called branched GDGTs (brGDGTs), which display characteristics typically found in archaeal membrane lipids, i.e., ether bonds and a membrane spanning nature (Schouten *et al*., 2013). Microbial membrane lipids can be used as taxonomic markers for specific microbial groups when found in the environment (Damsté *et al*., 2000; Hamersley *et al*., 2007; Rush and Sinninghe Damsté, 2017). Besides, microbial membrane lipids may change as a response to environmental changes to maintain the homeostasis of the membrane, thus also providing information on the physiological status of the cell (Sohlenkamp and Geiger, 2016). Lipids can be preserved in the sedimentary record longer than most other biomolecules like DNA or proteins, which makes them potential biomarkers for the presence of specific microbial groups or environmental conditions in the past (Brocks and Pearson, 2005; Rush and Sinninghe Damsté, 2017). Microbial (iso and br)GDGTs preserved in the sedimentary record are widely used as proxies to reconstruct past environmental conditions (Pearson and Ingalls, 2013; Schouten *et al*., 2013). Isoprenoid GDGTs are known to be synthesized by several archaeal phyla in soils, freshwater, and in the marine environment (Schouten *et al*., 2013; Fig S1). Branched GDGTs are found in soils, peats, and in freshwater systems (Pancost and Sinninghe Damsté, 2003; Weijers *et al*., 2006, 2007) and have also been detected in marine oxygen minimum zones (Liu *et al*., 2014) suggesting multiple bacterial source types and physiologies producing these lipids in the environment. Members of the bacterial phylum Acidobacteriota have been seen to synthesize the building blocks of brGDGTs (*i.e.*, iso-diabolic acid; Sinninghe Damsté *et al*., 2011) but in general the bacterial sources of brGDGTs are unknown. Recent genomic analyses based on the presence of biosynthetic genes suggest that multiple bacterial, and potentially even archaeal groups, might be able to synthesize brGDGTs (Sahonero-Canavesi *et al*., 2022).

To constrain the biological source(s) of GDGTs or any other lipid, the ultimate proof is the isolation of the producer and confirmation of the lipid production in the laboratory. However, only a small percentage of microorganisms have been cultured in the lab (Tyson and Banfield, 2005; Overmann *et al*., 2017; Lewis *et al*., 2021) thus preventing this approach in most cases. Alternatively, it is possible to investigate the genetic capacity of a microorganism to synthesize a lipid compound of interest in the environment without lab culture by detecting the genes coding for enzymes involved in the production of these lipids (Pearson *et al*., 2007; Welander *et al*., 2010). Most of the enzymes involved in the isoGDGT biosynthetic pathway are known (Koga and Morii, 2007; Jain *et al*., 2014); recently, several key steps in the pathway have been elucidated, such as the formation of the monolayer by coupling of the terminal-ends of the isoprenoids by the radical S-adenosyl-L-methionine (SAM) enzyme Tes (tetraether synthase; Lloyd *et al*., 2022; Zeng *et al*., 2022) as well as the formation of cyclopentane rings mediated by radical SAM GDGT ring synthases (gyrases GrsAB; Zeng *et al*., 2019). A recent study has also suggested the involvement of a radical SAM protein, a glycerol monoalkyl glycerol tetraether (GMGT) synthase (Gms), mediating the formation of a covalent bond between the two isoprenoid chains in isoGDGTs to form GMGT (previously known as H-GDGTs; Garcia *et al*., 2024). On the other hand, the lipid biosynthetic pathway of brGDGTs seems to be more diverse and is not yet fully constrained. Recently, we identified the MSL synthase (Mss) and glycerol ester reductase (Ger), involved, respectively, in the coupling of two FAs and the conversion of ester to alkyl ether bonds in brGDGTs under anaerobic conditions (Sahonero-Canavesi *et al*., 2022). Homologues of Tes that are found in bacterial genomes have also been proposed to be involved in the coupling of fatty acids in the formation of brGDGTs, but the activity of these enzymes has not been proven (Zeng *et al*., 2022). Besides, the enzymes encoded by the ether lipid biosynthesis gene cluster *elb* (i.e., *elb*B, D and E) in myxobacteria have been seen to be involved in the formation of alkyl ethers (Lorenzen *et al*., 2014), thus suggesting multiple potential ways to synthesize ether bonds in membrane lipids. Additionally, an alternative pathway involving an alkylglycerone phosphate synthase encoded by the *agps* gene is involved in the formation of alkenyl ethers (plasmalogens) in *Myxococcus xanthus* (Lorenzen *et al*., 2014) while alkenyl ethers can also be synthesized by the products of the *pls* operon under anaerobic conditions (Jackson *et al*., 2021). Even though specific microbial groups have been predicted to make iso and brGDGTs based on the presence of the above-mentioned genes in their genomes, constraining the producers in environmental samples is essential for making better interpretations of their use as biomarkers of paleoenvironmental proxies, their adaptive value to the microorganism, as well as to elucidate the timing of the evolutionary acquisition of these membrane lipids.

Here, we investigated the microbial sources of membrane spanning lipids (MSL, iso and brGDGTs) by using genome-resolved metagenomics targeting their lipid biosynthetic pathways in the Black Sea, the biggest euxinic (sulfidic and anoxic) basin in the world, where both iso and brGDGTs have been previously detected (Schouten *et al*., 2000; Liu *et al*., 2014).In addition, we report the brGDGTs composition and distribution in the Black Sea water column and compare them with microbial MSL potential producers to better constrain the sources of these lipids. Data derived from this study will also further aid in the cultivation and validation of the production of iso and brGDGTs of microorganisms present in the marine environment.

## Results and Discussion

In this study, we investigate the isoGDGT lipid data by Sollai *et al*., (2019) and we also report brGDGT lipid data obtained from the same samples of the Black Sea water column from 50 to 2,000 m depth—which were extracted and analyzed with a different approach (see Experimental procedures for details). In addition, DNA was extracted from the same samples and 16S rRNA gene amplicon sequencing, 16S rRNA gene quantification using quantitative PCR, and genome-resolved metagenomics (previously reported in Ding and von Meijenfeldt *et al*., 2024) was performed to determine the taxonomy, abundance, and lipid biosynthetic pathways of potential iso- and brGDGTs producers in this system.

### Physicochemical conditions, general microbial diversity and abundance in the Black Sea water column

Suspended particulate matter (SPM) samples had been collected from the Black Sea water column in high resolution from 50 to 2,000 m depth. Oxygen and sulfide concentrations were previously reported in Sollai *et al*., (2019) and summarized in Fig. S2. In brief, oxygen concentration decreased from 50 to 70 m depth, with suboxic waters from 70 to 110 m depth, and euxinic waters spanning down to 2,000 m depth where oxygen was below the detection limit and the sulfide concentration was approximately 400 µM (Fig. S2).

Microbial diversity was evaluated by 16S rRNA gene amplicon sequencing using universal primers to capture a broad phylogenetic diversity. The same 16S rRNA gene primers were used for estimating 16S rRNA gene absolute abundances attributed to different archaeal and bacterial groups based on quantitative PCR (see Experimental procedures for details).

Bacterial 16S rRNA gene reads were predominant throughout the water column, with values higher than 80% of the total (bacteria + archaea + unassigned at the domain rank) except for the depths between 250-1,000 m where bacterial relative abundance was lower than 70% (Table S1). Archaeal 16S rRNA gene reads ranged from 1 to 11% of the total throughout the depth profile, with maximum relative abundance in the suboxic zone (70-100 m), which then decreased to values around 2% in the euxinic zone (Table S1).

Bacterial diversity profiles based on 16S rRNA gene amplicon sequencing revealed a predominance of Bacteroidetes, Cyanobacteria, Proteobacteria and Verrucomicrobia in the oxic waters (i.e., 50 m depth) (Fig. 1, Table S2). In suboxic waters (i.e., 70-110 m depth), the 16S rRNA gene amplicon sequencing profiles indicated major presence of the phyla Actinobacteria (mainly class Acidimicrobiia, 12%), Bacteroidetes (mainly class Bacteroidia, up to 16% at 80 m), Epsilonbacteraeota (mainly class Campylobacteria, up to 30% at 170 m), Planctomycetota (up to 15%), and Marinomicrobia (up to 16%). Sequences attributed to the phylum Proteobacteria were also present in the suboxic waters, with the classes Alphaproteobacteria predominant in the upper suboxic waters (up to 15%), Gammaproteobacteria in the lower suboxic waters (up to 20%), and Deltaproteobacteria in the lower suboxic waters (up to 18%), also extending their presence in the euxinic waters (Fig. 1, Table S2). Some of the Deltaproteobacteria 16S rRNA gene sequences at these depths can be attributed to sulfate reducing bacteria (Table S2), which fits with a relevant role and presence of anaerobic organic matter processing sulfate reducers in the lower suboxic and euxinic waters of the Black Sea (van Vliet *et al*., 2021). Finally, bacterial 16S rRNA gene reads in the euxinic waters were dominated by members of the phylum Chloroflexi, (specifically classes Anaerolineae and Dehalococcoidia) increasing in abundance, from 130 meters downwards, -up to 30% of all reads; members of the phylum Cloacimonadota (prev. ‘*Candidatus* Cloacimonetes’) only present from 500 meters depth downwards and to a maximum of 14% between 1,000-2,000 meters (Fig. 1, Table S2). The here reported bacterial diversity coincides with previously reported studies in the Black Sea water column in different geographical stations, years and seasons (Cabello-Yeves *et al*., 2021; Pavlovska *et al*., 2021; Suominen *et al*., 2021), which suggests that the microbial population is highly stable, and that this system allows a comprehensive study with conclusions applicable to larger spatial and temporal scales.

**Figure 1.**
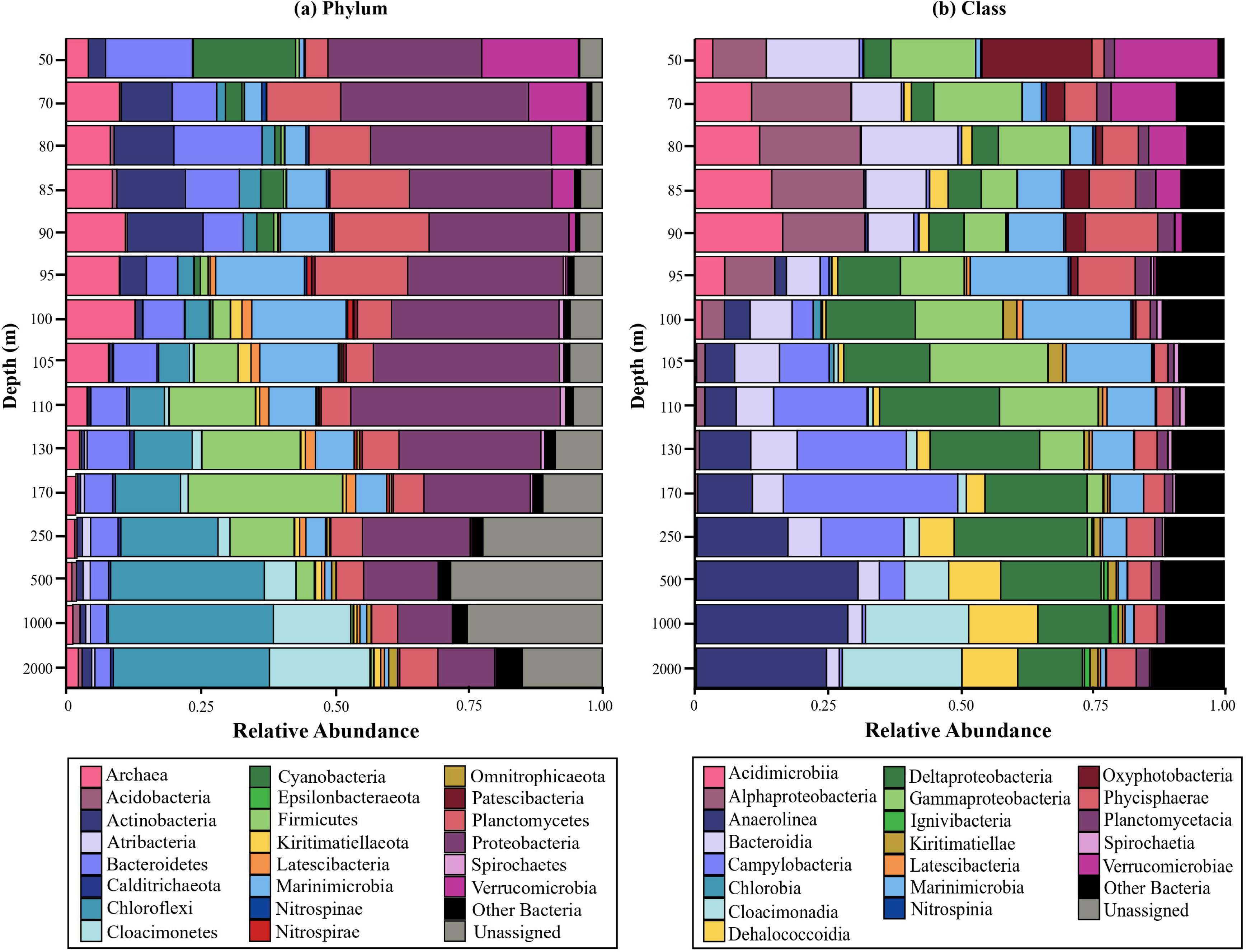
General bacterial diversity at the (a) phylum and (b) class rank based on 16S rRNA gene amplicon sequencing data in the Black Sea water column.

Archaeal diversity profiles based on 16S rRNA gene amplicon sequencing indicated that the phylum Thaumarchaeota (Nitrososphaerota), specifically the class Nitrososphaeria, constituted 85-90% of total archaeal 16S rRNA gene reads in suboxic waters and then the percentage declined downward (Fig. 2, Table S1). Members of the class Bathyarchaeia (phylum Crenarchaeota) contributed on average 40% of total archaeal 16S rRNA gene reads from 250-2,000 m, while class Thermoplasmata (phylum Euryarchaeota) were present in the entire water column, with those Thermoplasmata affiliated to Marine Euryarchaeota group II abundant in the oxic and suboxic waters, whereas those attributed to uncultured Thermoplasmatales mostly present in euxinic waters (Fig. 2A, Table S2). Sequences attributed to the class Woesearchaeia (phylum Nanoarchaeota, DPANN superphylum) contributed up to 40% of the total archaeal 16S rRNA gene reads in the upper euxinic waters but were also present in suboxic and euxinic waters (Fig. 2A, Table S2). In general, the archaeal diversity reported here by using 16S rRNA gene amplicon sequencing is similar to that previously reported in the same samples with different sequencing primers and a lower amount of sequenced reads by using another sequencing platform (Sollai *et al*., 2019).

**Figure 2.**
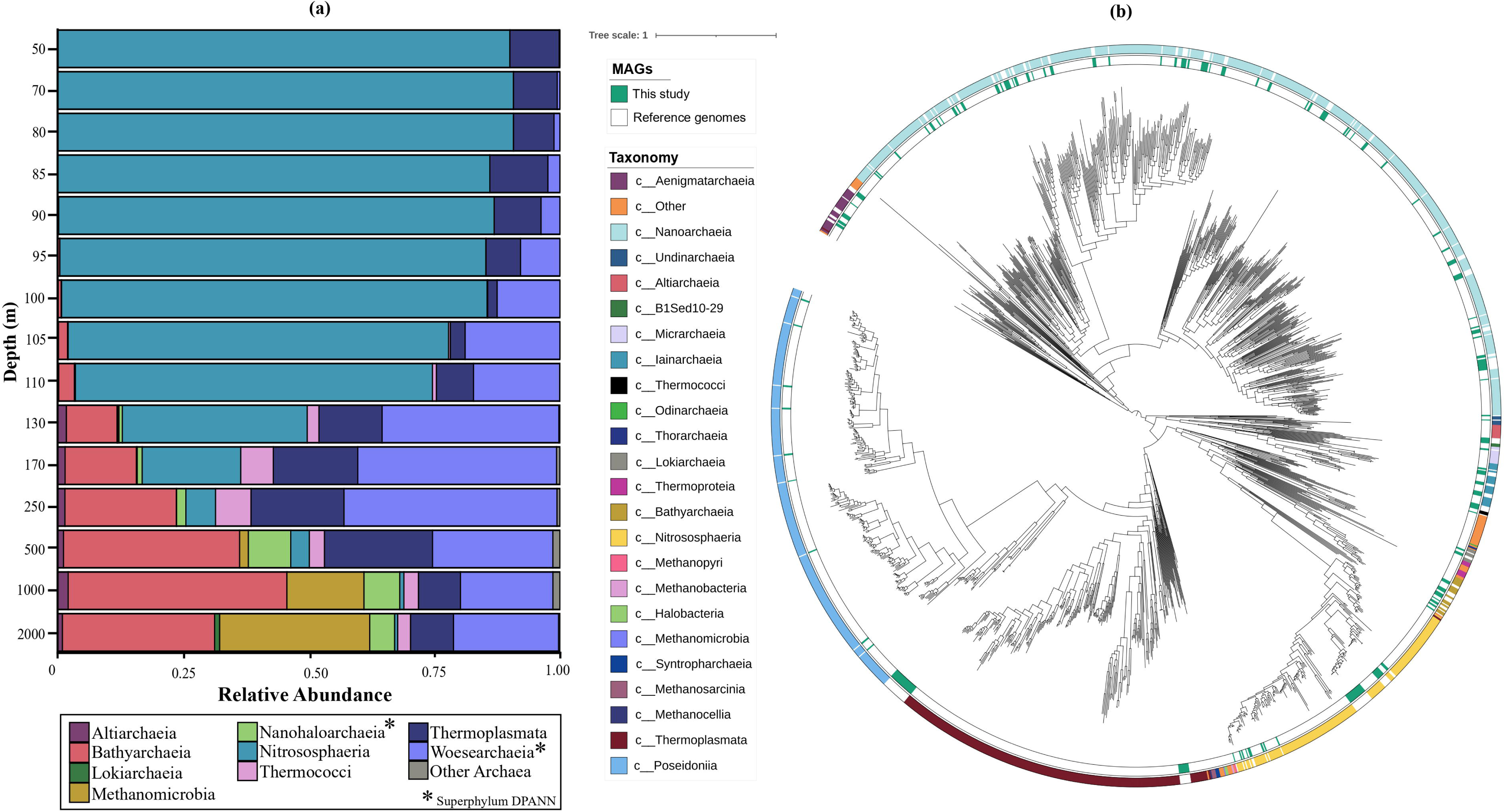
General archaeal diversity based on (a) 16S rRNA gene amplicon data (at the class rank), and (b) phylogenetic tree rooted at mid-point, with the metagenome-assembled genomes (MAGs) from the Black Sea (this study) in green and reference genomes in white, taxonomically classified according to the GTDB-Tk).

Nevertheless, due to deeper sequencing efforts, new archaeal groups were detected in the current study including members of the class Altiarchaeia (phylum Altiarchaeota) and of the order Aenigmarchaeales (class Nanohaloarchaei, part of the phylum Nanoarchaeaeota, DPANN superphylum) which accounted for approx. 2% and 7% of the total archaeal 16S rRNA reads in the euxinic waters, respectively (Table S2).

Total estimated archaeal abundance based on quantitative PCR showed a different distribution to that observed by Sollai *et al*., (2019) in the same samples, showing highest abundance of archaeal 16S rRNA gene reads at 85 m depth at the upper interface of the suboxic zone (Fig. S3, Table S3), while in the previous study the maximum abundance was detected well within the suboxic zone (70-100 m), showing the discrepancies that can be induced by using different 16S rRNA gene amplification primers on the same samples (e.g., Tremblay *et al*., 2015). Bacterial abundance estimated as total bacterial 16S rRNA gene copies per liter of SPM indicated maximum values at 85 m depth (Fig. S3, Table S3). Maximum bacterial abundance in the euxinic waters was detected at 1,000 m coinciding with the maximum of archaeal 16S rRNA gene copies L^-1^ but one order of magnitude higher than those attributed to archaea (Fig. S3, Table S3).

Archaeal diversity was further investigated by determining the diversity of metagenome-assembled genomes (MAGs) classified as Archaea that were obtained by sequencing the metagenome followed by de novo assembly and binning of the scaffolds of the same extracted DNA used for the amplicon sequencing (Ding and von Meijenfeldt *et al*., 2024; see Experimental procedures for details). MAGs affiliated to the phyla Aenigmatarchaeota, Altiarchaeota, Asgardarchaeota, Hadarchaeota, Ianarchaeota. Micrarchaeota, Nanoarchaeota (e.g. of orders Woesearchaeales and Pacearchaeales), Thermoplasmatota (of classes Poseidoniia and Thermoplasmata), Thermoproteota (of class Bathyarchaeia and order Nitrososphaerales), and Undinarchaeota were detected (classified based on GTDB-Tk, see Table S4). To further visualize this wide taxonomic diversity detected in the Black Sea, we generated a phylogenetic tree of the archaeal MAGs. MAGs were dereplicated and further selected on the basis of their completeness (≥50%) and used to build a phylogenetic tree with a selection of known archaeal genomes based on a concatenated alignment of 24 core vertically-transferred genes (Fig. 2B; see Experimental procedures for details). The archaeal taxonomic groups detected by metagenomic sequencing were largely similar to the groups detected in this study by 16S rRNA gene amplicon sequencing (Fig. 2A), indicating that the 16S rRNA gene primers used in this study are not considerably taxonomically biased. Some minor inconsistencies exist between the taxonomic profiles generated via amplicon sequencing and the taxonomic groups identified with metagenomic sequencing which arise because of the use of different taxonomic analysis pipelines. For example, Bathyarchaeia are considered part of the phylum Crenarchaeota in our 16S rRNA gene amplicon sequencing analysis, which is considered the phylum Thermoproteota in our metagenomes. For the remaining text, figures and tables, we will be using the CAT and BAT taxonomic classifications of the scaffolds and MAGs.

### Membrane lipid diversity in the Black Sea water column

#### Isoprenoid GDGTs diversity and distribution, and potential producers

Here, we used the isoGDGT data previously reported in Sollai *et al*., (2019) represented as the sum of the all intact polar lipid (IPL)-derived core lipids present in the water column at a determined depth. The distribution of archaeal lipids, i.e., archaeol and isoGDGTs, in the water column showed that archaeol was mostly present in deep euxinic waters, while isoGDGTs were abundant in the surface (50 m), suboxic, and in the deep euxinic waters (Fig. 3A, Table S5). Few archaeal lipids were present at 170-500 m depth (Fig. 3B). Although both isoGDGTs with no rings (i.e., GDGT-0) and with rings (GDGT-1-4 and crenarchaeol) had similar vertical distribution, the presence of isoGDGTs with rings increased with depth in comparison with the surface water where GDGT-0 was predominant (Fig. 3B).

**Figure 3.**
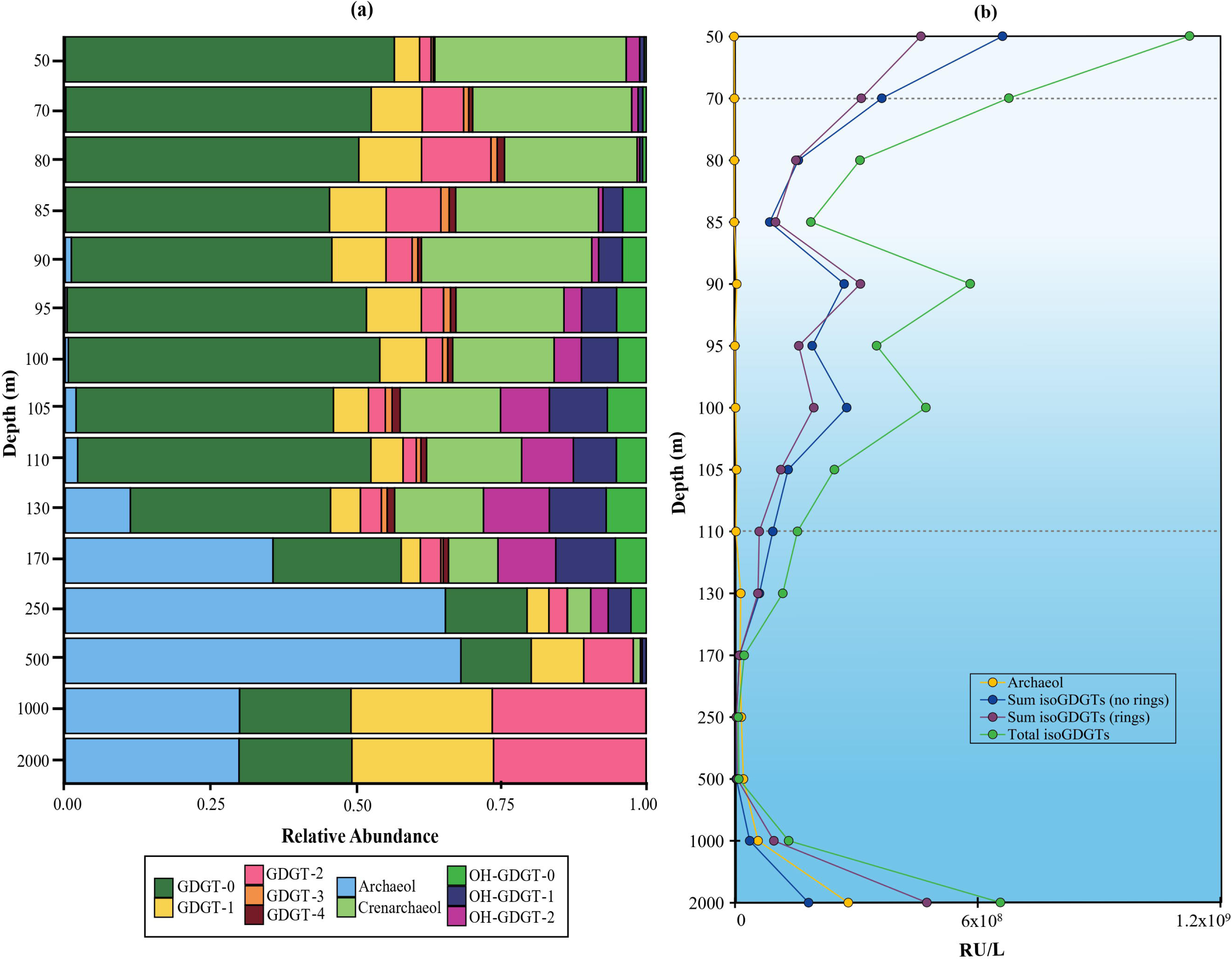
Distribution of archaeal isoGDGTs and archaeol (sum of intact polar lipid-derived core lipids reported by Sollai et al., 2019) in the Black Sea water column, (a) relative abundance, (b) response units (RU) per liter of isoGDGTs with no rings (*i.e.,* GDGT-0), with rings and total isoGDGTs. Dotted lines indicate the beginning of suboxic (70 m) and euxinic (110 m) zones.

Sollai *et al*., (2019) suggested that the lipid composition in the oxic and suboxic waters was likely mainly attributed to members of the Thaumarchaeota and of the Marine Euryarchaeota group II, whereas the lipids detected in the deep euxinic waters could be synthesized by archaea belonging to the DPANN superphylum, as well as classes Thermoplasmata, Bathyarchaeia, and members of ANME-1b, which were suggested to be responsible for the synthesis of archaeol, GDGT-0, GDGT-1, and GDGT-2. The observed GDGT-1 and 2 in the euxinic waters of the Black Sea have been traditionally thought to be produced in situ by members of the methanotrophic ANME group (e.g., Wakeham *et al*., 2003; Blumenberg *et al*., 2004). Although molecular carbon isotope evidence has supported the transfer of isotopically depleted methane-derived carbon into lipids of ANME archaea in the deep euxinic waters of the Black Sea, the ^13^C values measured in the bulk GDGT building blocks biphytanes have been previously reported to be substantially higher than observed in most other environments containing anaerobic methanotrophic archaea (Wakeham *et al*., 2003). An explanation for this offset might be additional or alternative sources to ANME-1 for isoGDGTs with rings in the deep euxinic waters of the Black Sea. The study by Sollai *et al*., (2019) showed that other archaeal groups like members of the order Thermoplasmatales (phylum Thermoplasmatota) and of the class Bathyarchaeia (phylum Thermoproteota) outnumbered ANME-1 at these depths, so potentially these groups could be also synthesizing isoGDGTs with rings and therefore invalidate the use of GDGT-1 and 2 as exclusive biomarkers of anaerobic methane oxidizers in similar marine systems. Among these archaeal groups, the presence of Bathyarchaeia and Thermoplasmatales in the deep euxinic waters is surprising, as these are archaeal groups normally found in sediments rather than in the water column. Their membrane lipid composition has never been assessed but previous studies have inferred the synthesis of GDGT-0 (Buckles *et al*., 2013) or butanetriol dialkyl glycerol tetraethers (BDGTs) (Meador *et al*., 2014) in members of the Bathyarchaeia based on the co-occurrence of lipids and these archaeal groups. However BDGTs have not been detected in these samples (Sollai *et al*., 2019; this study), suggesting these are not produced by the Bathyarchaeia subgroups present in the Black Sea water column at the time of sampling. The membrane lipid composition of yet-uncultured sedimentary Thermoplasmatales has never been assessed, therefore their potential of synthesizing GDGTs, with or without ring moieties, is uncertain. This is also applicable to the abundant archaeal DPANN superphylum (classes Nanohaloarchaeia and Woesearchaeia, Fig 2a) present in the euxinic waters (Sollai *et al*., 2019; this study). Nevertheless, some of the main DPANN groups present in the deep Black Sea are known to have very streamlined genomes lacking lipid biosynthetic capabilities (Castelle *et al*., 2018; Dombrowski *et al*., 2019, 2020), so it is likely they do not directly contribute to the GDGT pool in the Black Sea water column.

#### Branched GDGTs diversity, distribution, and potential producers

Previous studies have reported the presence of brGDGTs and overly-branched GDGTs with a higher degree of methylation (Weijers *et al*., 2006, 2007; De Jonge *et al*., 2015; Zeng *et al*., 2023) in the water column of oxygen minimum zones (Xie *et al*., 2014), in surface sediments (e.g., Liu *et al*., 2012; Xie *et al*., 2014; Becker *et al*., 2015) and lake sediments (e.g., Tyler *et al*., 2010; Niemann *et al*., 2012; Günther *et al*., 2014), as well as in the Black Sea water column (Schouten *et al*., 2000; Liu *et al*., 2014) which have been attributed to anaerobic planktonic microorganisms (Liu *et al*., 2014; Xie *et al*., 2014). An increase in the relative abundance of OB-GDGTs has been observed in sediments representing the Oceanic Anoxic Event 2, suggesting OB-GDGTs could be used to infer past marine anoxic conditions (Connock *et al*., 2022). To better assess the (paleo)proxy potential of these lipids, in-depth studies of their biological sources and their functional role are needed. In our study, both regular brGDGTs and OB-GDGTs were detected in the Black Sea water column as previously described (Liu *et al*., 2014), although they were outnumbered by the isoGDGTs from 130 m to 1,000 m depth (Fig. 4). Distribution of isoGDGTs, brGDGTs and OB-GDGTs was similar to that previously detected by Liu *et al*. (2014) in the Black Sea, with isoGDGTs being more abundant in the upper redoxcline, and brGDGTs and OB-GDGTs increasing in concentration with depth (Fig. 4). Nevertheless, the brGDGT distribution by Liu *et al*. (2014) reported a consistent increase of these lipids with depth up to 2,000 m, while in our samples, the distribution follows a less consistent pattern, with an increase observed until 130 m and then again at 1000 m and decrease both at 110 m and 500 m and subsequently at 2,000 m (Fig. 4).

**Figure 4.**
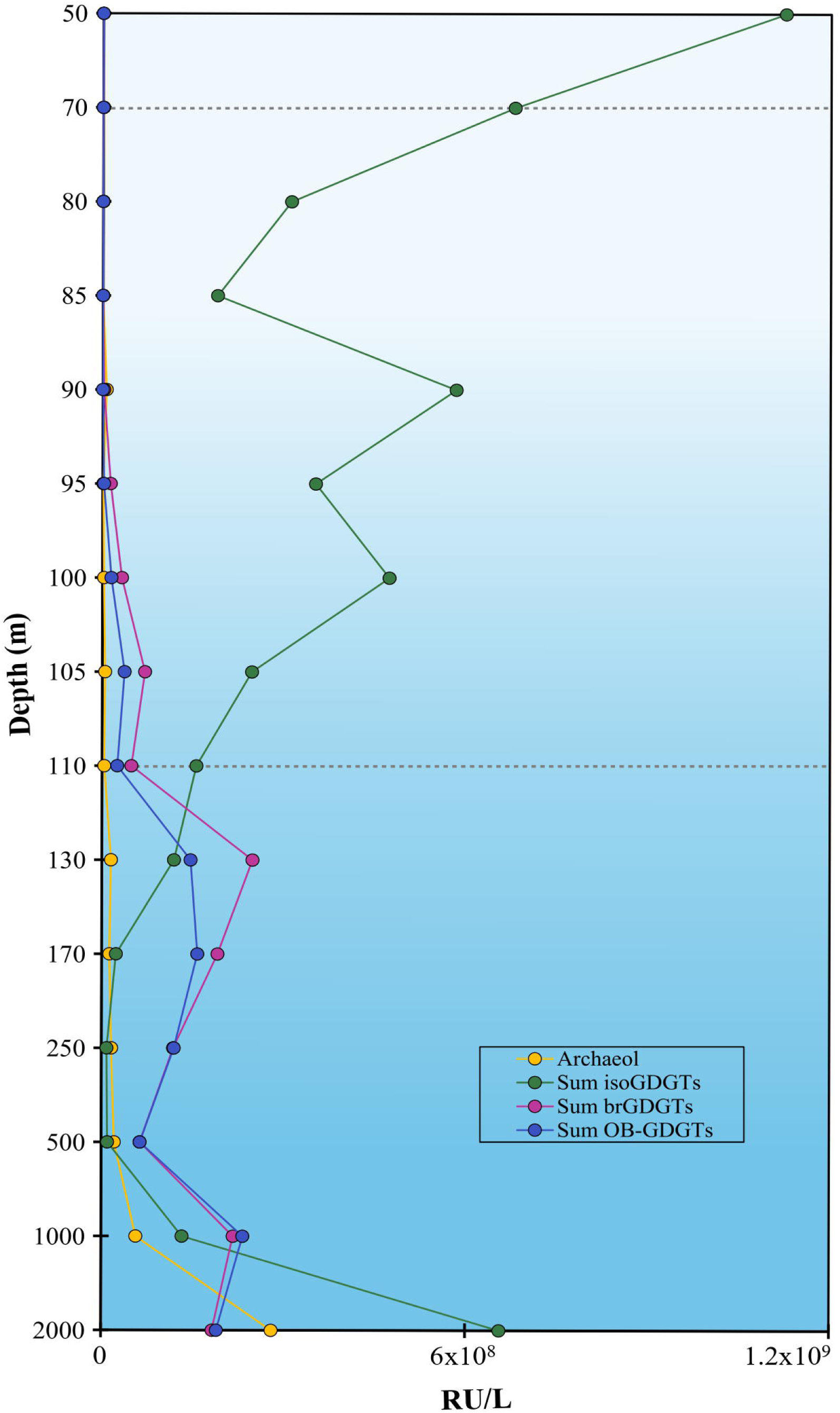
Distribution of the sum of isoGDGTs (intact polar lipid-derived core lipids), branched GDGTs (brGDGTs, core lipids) and overly branched GDGTs (OB-GDGTs, core lipids), as response units (RU) per liter in the Black Sea water column. Dotted lines indicate the beginning of suboxic (70 m) and euxinic (110 m) zones.

In general, previous studies and ours report a similar distribution of iso-, br- and OB-GDGTs in the water column of the Black Sea, reinforcing the idea that this euxinic system is very stable in terms of microbial diversity and producers of membrane lipids, over different years and seasons. In our study, we identified the brGDGTs and OB-GDGTs based on their accurate mass and elution patterns, however, for an easier visualization, we will refer the identified compounds by their nominal mass, both in the text and figures (for more details, see Table S7). We detected brGDGTs -1a (mass-to-charge, *m/z* 1022; Fig. S4A), -2a (*m/z* 1036, Fig. S4B), -2b (*m/z* 1034, Fig. S4D), and -3a (*m/z* 1050, Fig. S4C). Only brGDGTs -1a (*m/z* 1022), -2a (*m/z* 1036), and -3a (*m/z* 1050) were reported previously by Liu *et al*. (2014) in the Black Sea water column, with brGDGT-3a (*m/z* 1050) the most abundant from the suboxic waters downwards. The same distribution was observed in our study as brGDGT-3a (*m/z* 1050) was dominant in the euxinic waters followed by brGDGT-1a (*m/z* 1022) and -2a (*m/z* 1036) (Fig. 5A).

**Figure 5.**
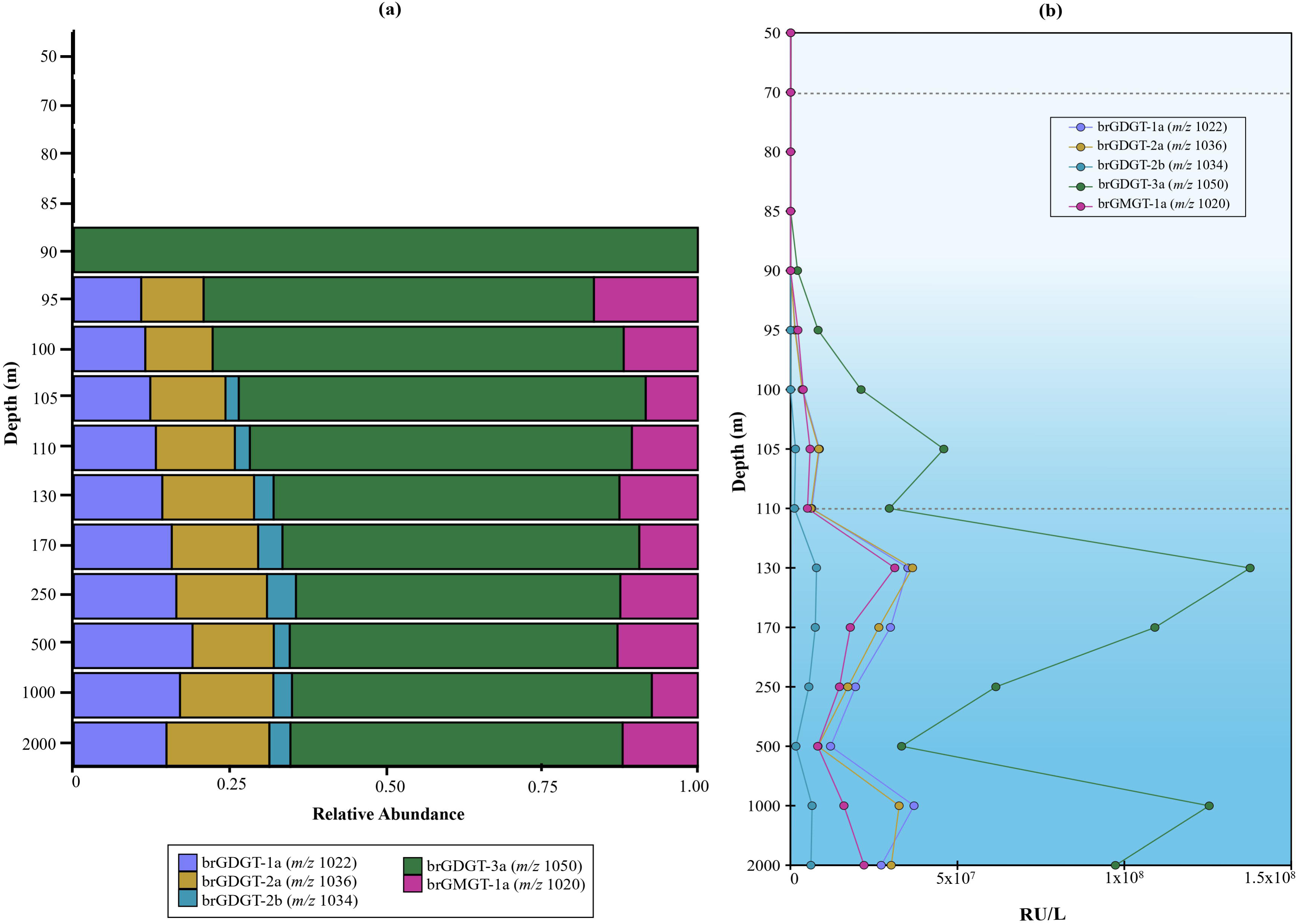
Distribution of branched GDGTs in the Black Sea water column, (a) relative abundance, (b) response units per liter. Dotted lines indicate the beginning of suboxic (70 m) and euxinic (110 m) zones.

The distribution of the detected brGDGTs as response units (RU) per liter indicated that all the detected brGDGTs had a similar distribution with local abundance maxima at 105, 130 and 1,000 m depth (Fig. 5B). Nevertheless, these detected abundance peaks were higher for brGDGT-3a (*m/z* 1050) than for the other brGDGTs, suggesting the potential contribution of additional producers of this brGDGT at that specific depth or specific environmental conditions that favor the production of brGDGT-3a.

We also detected the glycerol monoalkyl glycerol tetraether lipid (GMGT; also called ’H-GDGTs’) brGMGT-1a (*m/z* 1020) (Fig. S4E) with a maximum abundance at 130 m depth, followed by a decrease with depth, and then an increase from 1,000 to 2,000 m depth This was the only brGDGT that was higher in abundance at 2,000 m than 1,000 m depth (Fig. 5B).

Oxygen limitation is likely to trigger brGDGT production (Halamka *et al*., 2021), in terrestrial environments branched GMGTs are generally present at sites characterized by high nutrient levels and/or oxygen-limited conditions, suggesting anaerobic microbes as likely sources (Naafs *et al*., 2018; Elling *et al*., 2023). In the marine realm, brGMGT-1a (*m/z* 1020) has been detected in marine surface sediments (Liu, Lipp, *et al*., 2012), in suspended particulate matter from the oxygen minimum zone of the eastern Pacific (Xie *et al*., 2014), and in the Bay of Bengal (Kirkels *et al*., 2022), suggesting that anaerobic planktonic microorganisms are responsible for their production, in line with the presence of brGMGT-1a (*m/z* 1020) in the euxinic waters of the Black Sea.

We also detected a diversity of OB-GDGTs: OB-GDGT-9 (*m/z* 1092), OB-GDGT-10 (*m/z* 1106), OB-GDGT-11 (*m/z* 1120) and OB-GDGT-12 (*m/z* 1134) (see Fig. 6A, Fig. S4F-I, Table S6), which had been previously detected in the Black Sea water column by Liu *et al*. (2014). The distribution of the OB-GDGTs is similar to those detected by Liu et al. (2014), with the only exception, that the concentration of the regular brGDGTs and OB-GDGTs in our dataset decreases at 2,000 m, while Liu *et al*. (2014) reports a consistent increase with depth (Fig. 6B), as also seen for the brGDGTs.

**Figure 6.**
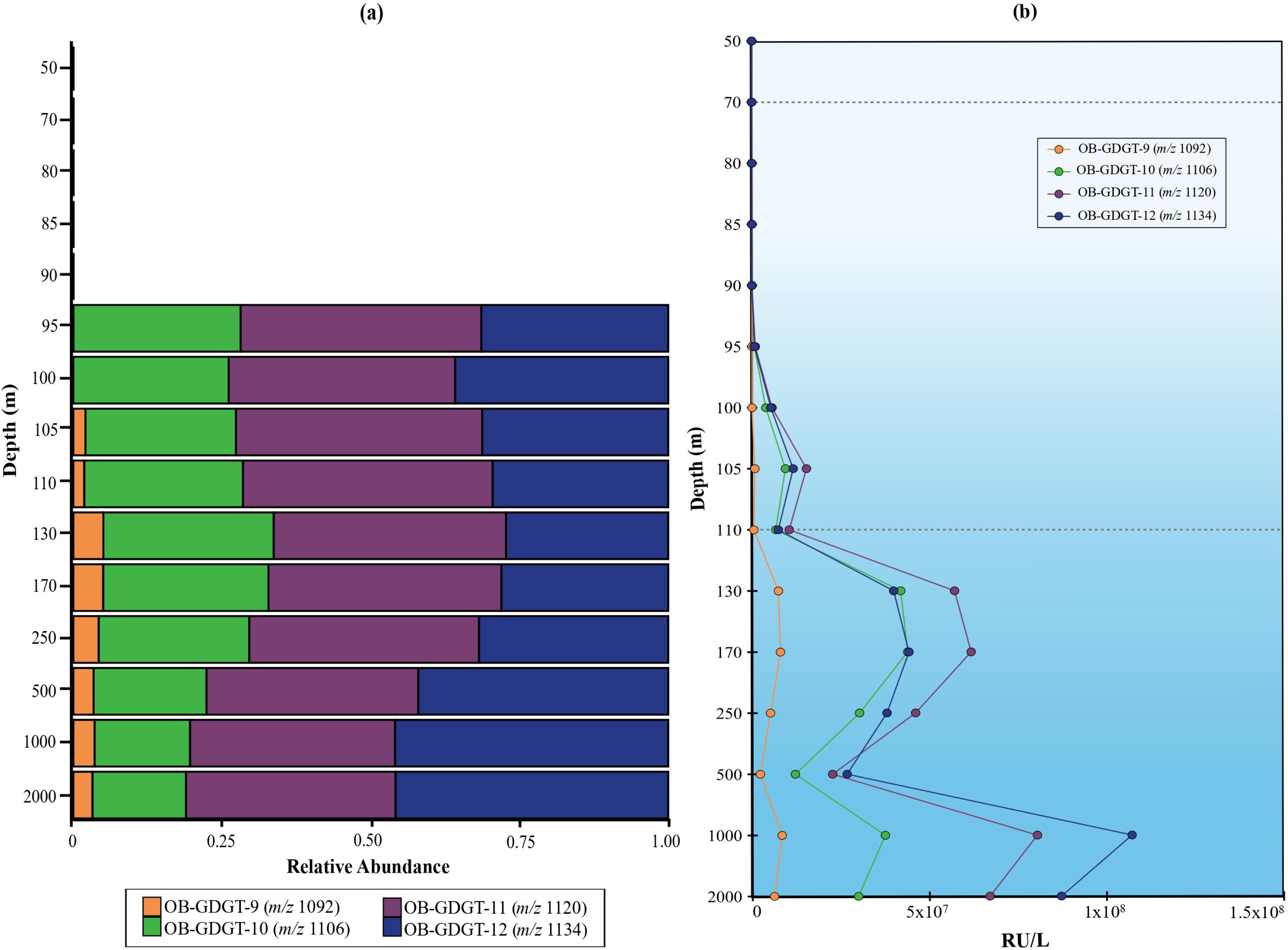
Distribution of overly branched OB-GDGTs in the Black Sea water column, (a) relative abundance, (b) response units per liter. Dotted lines indicate the beginning of suboxic (70 m) and euxinic (110 m) zones.

Zeng *et al*., (2023) investigated OB-GDGTs in sediments and observed a higher degree of methylation in the OB-GDGTs with increasing depth (higher % of OB-GDGT-9 (*m/z* 1092), OB-GDGT-10 (*m/z* 1106), and OB-GDGT-12 (*m/z* 1134) in deeper sediment depths. A similar observation was encountered in the marine water columns of the Black Sea and the Cariaco Basin (Liu *et al*., 2014).

Nevertheless, in our study, the detected OB-GDGTs did not change in relative abundance compared to each other with depth (Fig. 6B), thus suggesting that the production of branched and OB-GDGTs in the deep waters of the Black Sea is not consistent between different sampling sites and seasons, and may be associated with different producers or differences in the abundance of the existing populations at the deeper Black Sea waters.

Regarding the potential sources of brGDGTs and OB-GDGTs in marine anoxic systems, Zeng *et al*., (2023) performed a co-occurrence network analysis correlating the relative abundance of those lipids with that of the bacterial communities based on 16S rRNA gene amplicon sequencing. They suggested that members of the bacteria Chloroflexi, Proteobacteria and Dadabacteria could be the potential producers of brGDGTs and that Armatimonadota, Planctomycetota and Chloroflexi could be the producers of OB-GDGTs in the Mariana Trench sediments, pending further confirmation.

#### Inference of iso- and brGDGTs producers based on their lipid biosynthetic genomic potential

As mentioned above, previous studies have inferred the biological sources of iso-, br- and OB- GDGTs by correlating the presence of lipids in environmental samples with the occurrence of archaeal and bacterial groups based on 16S rRNA gene amplicon sequencing (Buckles *et al*., 2013; Sollai *et al*., 2019; Baxter *et al*., 2021; De Jonge *et al*., 2021; Zeng *et al*., 2023). However, this inference does not necessarily correctly link a product (lipid) with its sources as correlation does not imply causation (Ding and von Meijenfeldt *et al*., 2024). To make a more accurate link between a lipid and its producer(s), here we targeted the genomic capacity to synthesize these lipids in the assemblies of the sequenced metagenomes of the 15 samples throughout the Black Sea water column (∼1.64 x 10^8^ scaffolds and 5,844 metagenome-derived genomes (MAGs)) (see Experimental procedures for details), and compared the distribution of the hits with the distribution of the observed membrane lipids.

We performed two complementary analyses. First, we examined the fraction of MAGs from a certain phylum in each sample that encode a relevant key gene to synthesize the lipid of interest, which reflects the within-phylum strain diversity and the universality of the biosynthetic capacity in the phylum at different depths in the water column (Fig. 7, S10). This shows niche-based differentiation of different strains within a phylum, for example when all MAGs of a phylum in the upper water column have the biosynthetic capacity to synthesize a lipid and few in the deep waters do. In addition, this analysis allows for an assessment of noise due to technical artifacts, as it is likely that a high fraction of MAGs encoding a gene points to its true presence within the phylum instead of a false positive due to spuriously binned genes, especially if they represent a large number of MAGs (large circles and dark colors in Fig. 7, S10). Second, we quantified the abundance of hits in the assembly irrespective of whether the scaffold was binned or not, using normalized depth based on read mapping to the scaffolds (i.e., number of mapped reads per base pair per 1x10^8^ mapped reads, see Experimental Procedures for details) which allows for a comparison of sequenced DNA between samples (Fig. 8, 9, S5-9). This second analysis takes abundance into account, and allows for a correlation with the lipid profile. Moreover, since also unbinned scaffolds are examined, it represents a more comprehensive picture of the microbiome than the MAGs alone (Hauptfeld *et al*., 2024), even if the taxonomic annotations of scaffolds from highly unknown microorganisms are less robust than those of MAGs (von Meijenfeldt *et al*., 2019). For the case of the genomic capacity to synthesize isoGDGTs, here we determined the distribution and abundance of the detected homologs (see Experimental procedures for details) of Tes (GDGT synthase; Zeng *et al*., 2022) involved in the coupling of isoprenoids leading to GDGT without and with cyclopentane rings, and of the gyrase GrsA involved in the GDGT cyclization up to 4 rings (Zeng *et al*., 2019). Tes gene hits were detected in different microbial groups throughout the water column, within Archaea mostly in the phylum Thermoproteota (classes Nitrosphaeria and Bathyarchaeaia) and class Thermoplasmata (Fig. 7A, 8A, Table S8, S9AB). The distribution of isoGDGTs (Fig. 3B,4) can be compared to the abundance profile of Tes hits (Fig 8A), with those of the Thaumarchaeota genus *Nitrosopelagicus* more abundant in the surface waters (i.e., 70 m), followed by a maximum of Tes hits attributed to the Thaumarchaeota genus *Nitrosopumilus* in the upper suboxic waters (i.e., 100 m), while Tes hits of Bathyarchaeia and Thermoplasmata were more abundant in the upper euxinic waters (250 m), followed by an increase with depth of Tes hits of members of the ANME-1 Archaea (class Syntropharchaeia) (Fig. 8A, Table S8, S9AB).

**Figure 7.**
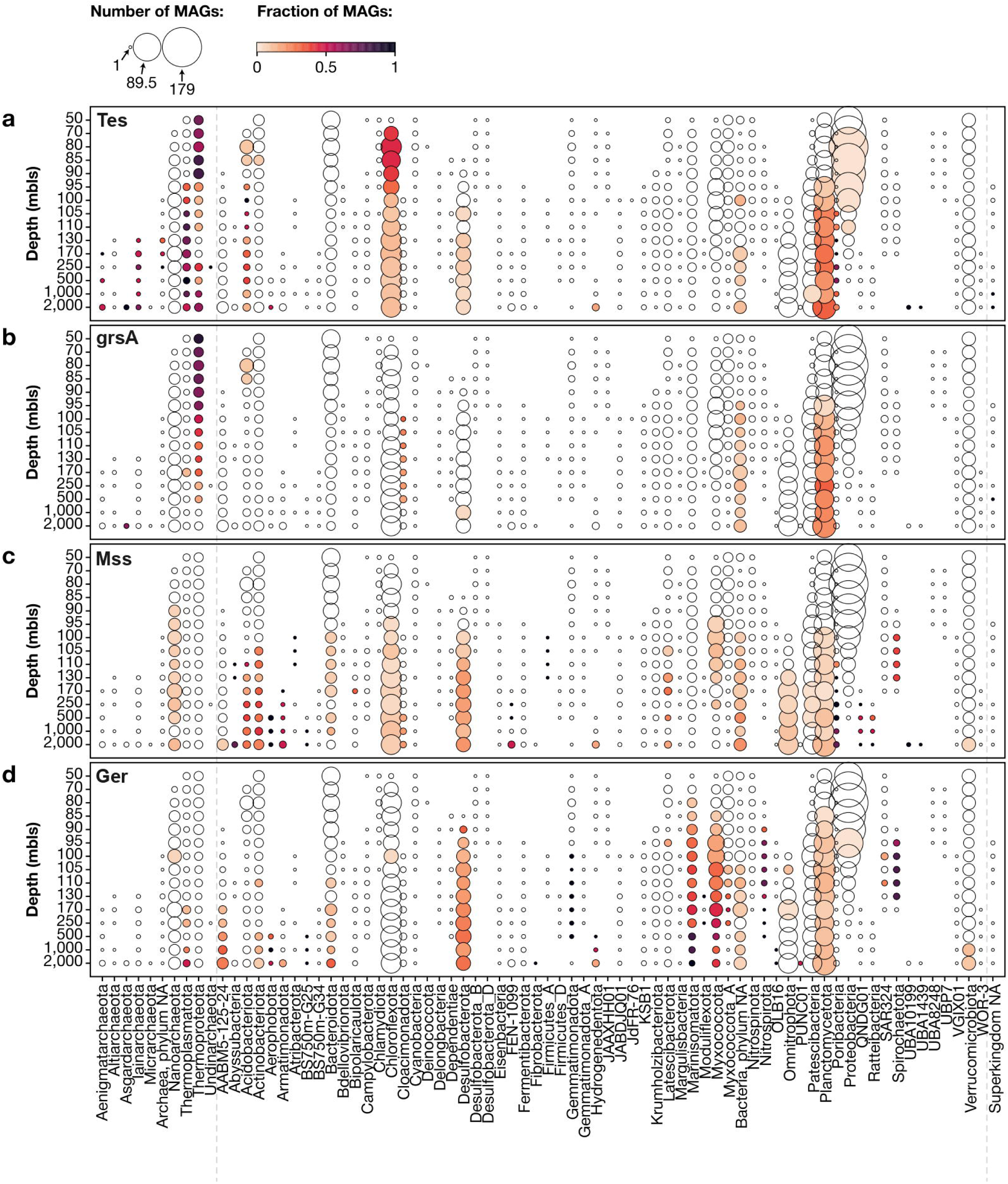
Number of metagenome-assembled genomes (MAGs) binned in the Black Sea, grouped by sample (depth) and taxonomy (rank: phylum). Circle size represents the number of MAGs by sample/taxonomy, color intensity describes fraction of MAGs containing protein homolog: a)Tes; b)GrsA; c) Mss; d)Ger.

**Figure 8.**
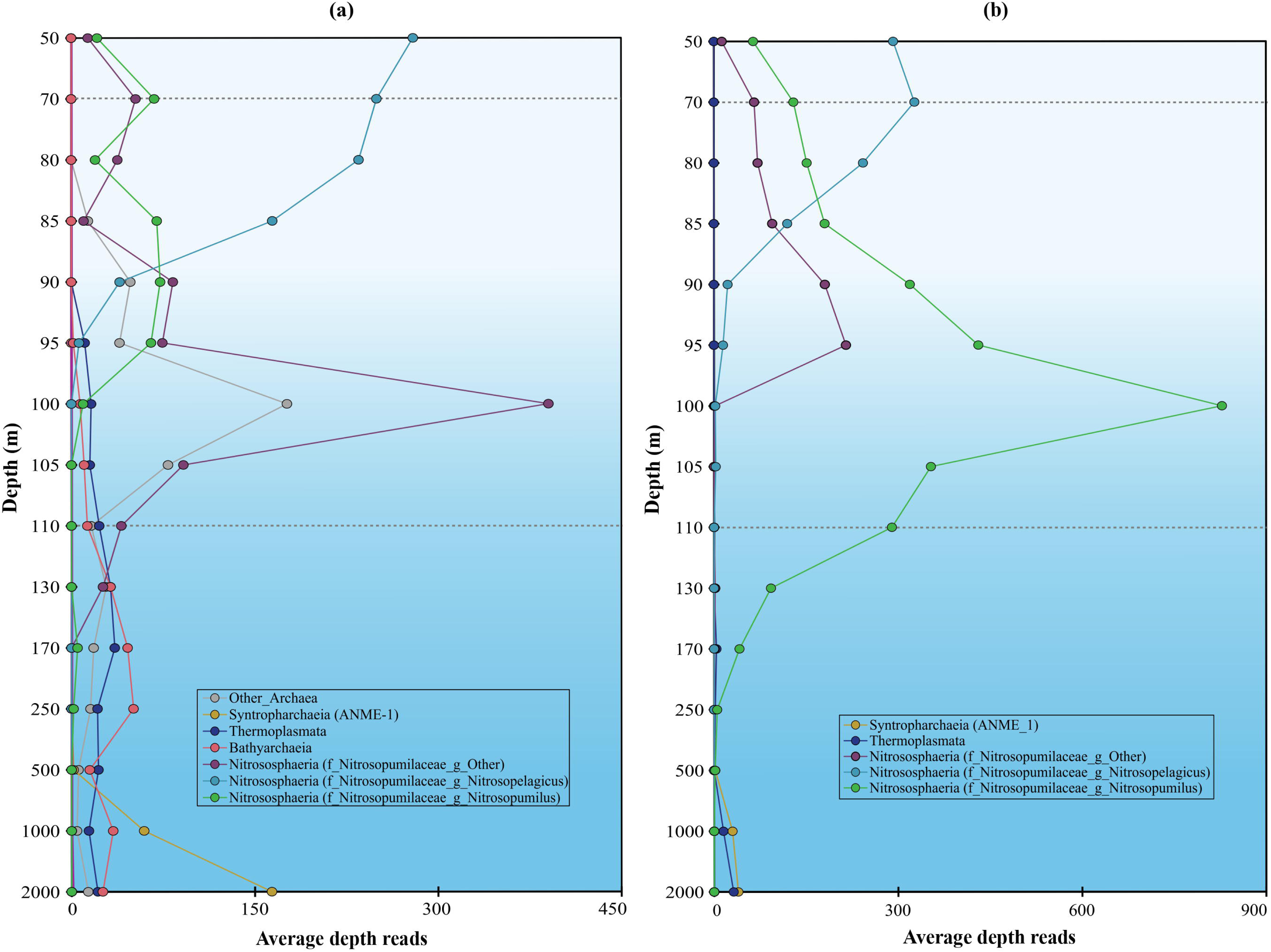
Sum of the average depth (i.e., number of mapped reads per base pair, per 1e+8 mapped reads) of Tes protein homolog hits (a) and of GrsA protein homolog hits (b) (see Experimental procedures for details) detected in archaeal groups across the Black Sea SPM profile from 50 to 2,000 m depth. (taxa marked with * have been plotted on the secondary axis for a better visualization). Dotted lines indicate the beginning of suboxic (70 m) and euxinic (110 m) zones.

**Figure 9.**
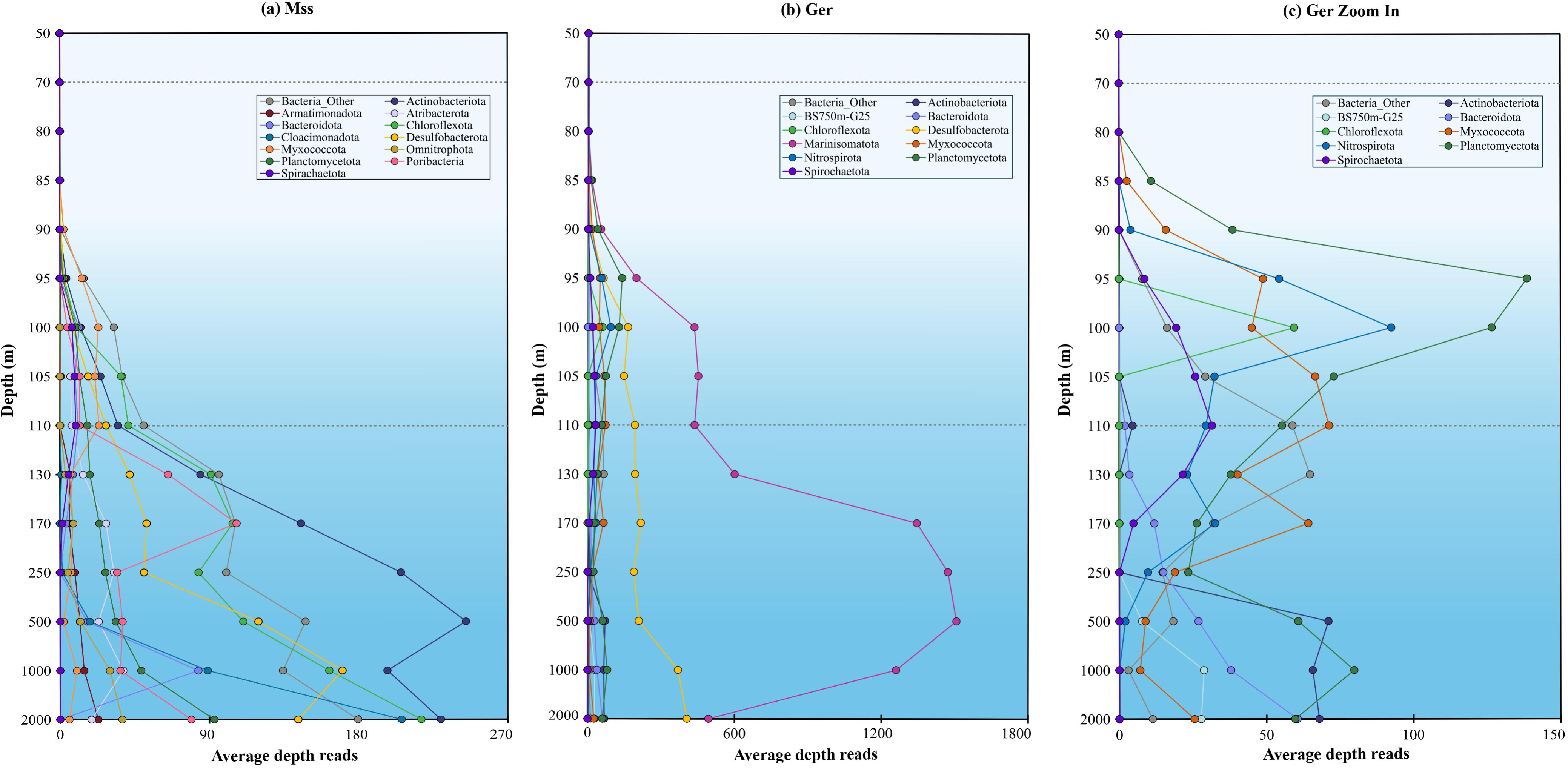
Sum of the average depth (i.e., number of mapped reads per base pair, per 1e+8 mapped reads) of the Mss protein homolog hits (a) and of the Ger protein homolog hits (b) with detail of less abundant taxa (c) (see Experimental procedures for details) detected in archaeal groups across the Black Sea SPM profile from 50 to 2,000 m depth. Dotted lines indicate the beginning of suboxic (70 m) and euxinic (110 m) zones.

This profile coincides with the distribution of hits of the gyrase GrsA for Thaumarchaeota (equivalent to GTDB-Tk classification Thermoproteota) genera *Nitrosopelagicus* and *Nitrosopumilus* (Fig. 8B, Table S8, S10AB), indicating that those two genera are likely the main and only producers of isoGDGTs with rings in the surface and suboxic waters. In the euxinic waters, we detected GrsA hits in low abundance attributed to ANME-1 (class Syntropharchaeia; 3 unbinned scaffolds from the 3 deepest samples), as well as to class Thermoplasmata (3 unbinned scaffolds from 250 m depth, 1,000 m depth, and 2,000 m depth, and 1 binned scaffold from 170 m depth (Fig. 8B, Table S10AB), suggesting these two archaeal groups could be responsible for the biosynthesis of isoGDGTs with rings in the deep waters of the Black Sea. However, evidence supporting this is weak as the scaffold where those GrsA homologs were detected are unbinned and their abundance is low. Is worth noting that members of the Thermoplasmata have been widely detected in marine sediments and anoxic water columns often co-occurring with ANME-1 (Blumenberg *et al*., 2004; Suominen *et al*., 2021; Iasakov *et al*., 2022; Wegener *et al*., 2022). If members of the Thermoplasmata present in similar settings would actually harbor the capacity to make isoGDGT-1 and 2, it would compromised the use of these isoGDGTs as markers for anaerobic methane oxidation by Archaea.

Tes homologs were also detected in 281 bacterial MAGs (Fig. 7A, Table S8). The presence of homologs of this gene in bacterial genomes was also observed in a microbial genome screening by Zeng *et al*., (2022) and could be indicative of their potential to synthesize brGDGTs. Although the presence of Tes homologs in bacterial genomes has not been experimentally confirmed to provide the capacity to synthesize bacterial membrane spanning lipids, here we report the distribution of Tes hits in bacterial genomes (Table S9AB), with members of the phylum Chloroflexota more abundant in the surface and upper suboxic waters, while others were more abundant in the lower suboxic and upper euxinic waters such as phyla Acidobacteriota, Desulfobacterota, Poribacteria and Planctomycetota, with the latter dominant in the suboxic and euxinic waters with a maximum at 2,000 meters depth (Fig. S5A). The bacterial diversity of Tes homologs detected in the MAGs (Fig. 7A) includes comparable taxonomic groups, with the most diverse groups being phyla Chloroflexota (Tes homologs detected in 111 out of 684 MAGs) and Planctomycetota (110 out of 684 MAGs), followed by Acidobacteriota (17 out of 97 MAGs), Desulfobacterota (10 out of 262 MAGs) and Poribacteria (10 out of 29 MAGs) (Table S8).

Hits of the gyrase GrsA enzyme were mostly detected in bacterial MAGs of phyla Planctomycetota (94 out of 684 MAGs) and Cloacimonadota (7 out of 42 MAGs) and was noticeably missing from members of the phylum Chloroflexota (Fig. 7B, S5B, Table S8, S10AB). Some additional hits were detected in Acidobacteria and Desulfobacteriota (Fig. S5B). We did not detect any hits of Tes (in MAGs and unbinned scaffolds) in phylum Cloacimonadota, possibly suggesting that the GrsA homologs in the phylum have a different function than GDGT cyclization or that Cloacimonadota encode a non-homologous gene that replaces the putative Tes-like functionality (note that many of the Cloacimonadota MAGs also do not encode Mss, see in Fig. 7C). Although the functional roles of Tes and GrsA have not been assessed in bacterial genomes, members of the Planctomycetota could potentially synthesize some of the brGDGTs with rings reported in our study (Fig. 5).

In addition, Black Sea MAGs were screened for the presence of proteins involved in the confirmed synthesis of bacterial membrane spanning lipids, namely Mss and Ger enzymes involved in the coupling of fatty acids and of the ether bond formation of brGDGTs, respectively (Sahonero-Canavesi *et al*., 2022). Mss hits (Tables S11AB) were abundant and detected in both unbinned scaffolds and MAGs of several bacterial phyla (Table S8). The abundance profile based on unbinned scaffolds (Fig. 9A), shows phylum Actinobacteriota as the most abundant microbial group, which increased with depth from 110 m downwards with maximum abundances at 500 m; phyla Chloroflexota, Cloacimonadota and Planctomycetota with a maximum abundance at 2000 m; and phyla Desulfobacterota and Bacteroidota with a maximum abundance at 1000 m (Fig. 9A). The MAGs diversity profile (Fig. 7C) includes the same taxonomic groups: Actinobacteriota (Mss homologs detected in 17 out of 146 MAGs), Chloroflexota (25 / 558 MAGs), Cloacimonadota (3 / 42 MAGs) and Planctomycetota (27 / 684 MAGs), Desulfobacterota (34 / 262 MAGs) and Bacteroidota (7 / 273 MAGs) with Mss protein-coding sequences additionally detected in MAGs of the phyla Acidobacteria (11 / 97 MAGs), Armatimonadota (7 / 13 MAGs), Latescibacteriota (7 / 92 MAGs), Myxococcota (12 / 347 MAGs), Omnitrophota (15 / 267 MAGs), Patescibacteria (3 / 428 MAGs), Poribacteria (14 / 29 MAGs), Spirochaeota (8 / 28 MAGs), among others (Fig 7C, Table S8). This diverse group of bacterial phyla hints at a more widespread potential to synthesize membrane spanning lipids in marine bacterial communities than previously considered. Although we did not detect any Tes hits in Cloacimonadota, the presence of hits for both GrsA (7 / 42 MAGs) and Mss (3 / 42 MAGs) could hint to the potential of this group to produce brGDGTs with rings, although co-occurrence of both genes in the same genome is inconclusive. Only 1 of the 42 surveyed MAGs has both enzymes, and this could be explained by either contamination, incomplete assembly and/or binning, or specific environmental conditions under which some of the organisms of this phylum could produce MSL using both enzymes.

Homologs of Ger (glycerol ester reductase) were detected in a diverse group of bacteria. The phylum Marinisomatota were amongst the most abundant in suboxic and euxinic waters (Fig 9B). Based on the unbinned scaffolds (Table S12AB), order UBA8477 was distributed towards the upper waters and the order Marinisomatales had a maximum abundance at 500 m. Marinisomatota was also amongst the most diverse groups (Ger homologs detected in 45 out of 134 MAGs) (Fig 7D, Table S8) . Other abundant (Fig. 9C) and diverse (Fig 7D, Table S8) phyla were the Myxococcota (61 / 347 MAGs), Desulfobacterota (54 / 262 MAGs), Planctomycetota (53 / 684 MAGs), Spirochaeota (19 / 28 MAGs), Nitrospirota (14 / 23 MAGs), Bacteroidota (9 / 273 MAGs), Actinobacteriota (4 / 146 MAGs), Aerophobota (4 /7 MAGs) (Table S8, S12AB).

Genomic coincidence of Mss and Ger hits were found in 34 MAGs of Desulfobacterota, 27 MAGs of Planctomycetota, 12 MAGs of Myxococcota, and 4 MAGs of Actinobacteria, pointing to their genomic capacity to synthesize brGDGTs and suggesting these might be the bacterial groups responsible for producing brGDGTs and OB-GDGTs detected in the Black Sea water column in this study. Note that Ger is a close homolog of PlsA which is an enzyme involved in the production of alkenyl ethers (i.e., plasmalogens). It is currently not possible to discriminate if the presence of these homologs leads to the formation of alkyl or alkenyl ethers based on sequence homology (Sahonero-Canavesi *et al*., 2022), and the distribution of Ger-like hits in different bacterial MAGs can also indicate the potential to produce plasmalogens in some of these groups, which were not part of this study.

Homologs of genes encoding the ElbD and Agps proteins of *M. xanthus* were also screened in the metagenomes to constrain the presence and abundance across the water column in different taxonomic groups which might harbor the capacity to synthesize alkyl or alkenyl ether bonds, respectively. For the case of ElbD, hits were found in a diverse group of bacterial phyla (Fig S10A Table S8). The most abundant (Fig. S6, Table S13AB) groups in the water column were the phyla Myxococcota, distributed predominantly in the upper oxic and suboxic waters; Chloroflexota abundant in upper suboxic and euxinic waters (peaks at 85, 500 and 2000 m); Proteobacteria in the upper suboxic waters; Actinobacteria, Omnitrophota and the candidate division AABM5-125-24 of the FCB superphylum in the euxinic waters (Fig. S6). These phyla were also amongst the most diverse (Fig S10A Table S8): Myxococcota (121 / 347 MAGs), Omnitrophota (93 / 267 MAGs), division AABM5-125-24 (58 / 69 MAGs), Proteobacteria (45 / 932 MAGs), Chloroflexota (37 / 558 MAGs); although other phyla containing many MAGs that encode elbD homologs (Fig. S10A) were found: Latescibacterota (50 / 92 MAGs), Verrucomicrobia (43 / 270 MAGs) and Planctomycetota (33 / 267 MAGs). Homologs to the gene encoding the Agps protein involved in the formation of alkenyl ethers in *Myxococcus* were also found in a wide diversity of bacterial MAGs binned from the Black Sea water column metagenomes (Fig. S10B, Table S8). Hits with the highest abundance across the water column were affiliated with phyla Actinobacteria, Proteobacteria and Verrucomicrobiota in the suboxic waters and with phyla Marinisomatota, Desulfobacterota and Chloroflexota in euxinic waters (Fig. S7). These phyla also corresponded to the most diverse: Desulfobacterota (Agps homologs detected in 59 / 262 MAGs), Chloroflexota (56 / 558 MAGs), Marinisomatota (49 / 134 MAGs), Proteobacteria (42 / 932 MAGs) and Verrucomicrobiota (43 / 270 MAGs) (Fig. S10B). Although not very abundant, Myxococcota (120 / 347 MAGs) and Latescibacterota (67 / 92 MAGs) and Myxococota were amongst the most diverse phyla encoding *agps* homologs (Fig. S10B).

Our metagenomic screening supports that the genomic capacity to synthesize alkyl ethers via Ger or ElbD, and alkenyl ethers via Agps is harbored by a wide diversity of bacterial groups which seem to be distributed in different niches throughout the Black Sea water column. MAGs of the phylum Verrucomicrobia, specifically of the *Pontiella* genus, were found to contain homologs coding for the ElbD protein. Members of the *Pontiella* genus have been isolated in a previous study from Black Sea samples and their lipids have been analyzed, but no ether-bound lipids were detected (Van Vliet *et al*., 2020). This emphasizes that the genomic presence of homologs of this gene does not necessarily imply the synthesis of the lipids, either because the gene is not involved in the lipid’s biosynthesis or because it is not expressed under laboratory conditions. MAGs affiliated to Acidobacteria containing *elbD* homologs (15 / 97 MAGs) were also found in the Black Sea water column present throughout suboxic and euxinic waters (Table S8, S13AB). Some members of the Acidobacteria are known to synthesize brGDGTs (Sinninghe Damsté *et al*., 2014, 2018), but the pathway used for their biosynthesis is still unclear as only some genomes of Acidobacteria subdivision 4 harbor the *elb* gene cluster or *agps* homologs and the role of the proteins that they encode in the ether bond formation has not been conclusively demonstrated. Therefore, it is unclear if the uncultured Acidobacteria detected in the Black Sea could be potential sources of the detected brGDGTs; however, it is worth mentioning that 9 MAGs of the phylum Actinobacteriota (class Humimicrobiia) obtained from our dataset have been seen to harbor homologs of both the Mss (Table S8, S11AB) and of the ElbD proteins (Table S8, S13AB), indicating genomic potential of members of this group to synthesize brGDGTs using the *elb* gene cluster to make alkyl ether bonds rather than by using the Ger protein. In this regard, out of the 558 MAGs from the phylum Chloroflexota present in the Black Sea water column, only 25 MAGs were detected with Mss and 37 MAGs with ElbD, and from these, a small fraction (9 MAGs) contains homologs of both protein-coding genes, suggesting different potential genomic capacities to make brGDGTs within the same phylum (Tables S11AB-S13AB). The same applies to MAGs of members of the phylum Myxococcota (12 MAGs with Mss homologs, 61 MAGs with Ger homologs, 121 MAGs with ElbD homologs, 120 MAGs with Agps homologs, and a smaller subset of MAGs with two or more of the homologs) . The phylum Planctomycetota represents an even stranger case. Homologs of Tes and GrsA (previously thought to be exclusive to Archaea) were detected in some representatives of the class Phycisphaerae of this phylum, while Mss and Ger homologs were detected in the Brocadiia, Phycisphaerae, and Planctomycetia classes. This suggests that members of the Planctomycetota phylum can be a source of very diverse lipids, including archaeal-like lipids with rings or brGDGTs. Some members of Brocadiia have been seen to produce alkyl ethers (Rattray *et al*., 2008) which in some cases coincides with the presence of either Ger, ElbD or Agps homologs in their genome (Sahonero-Canavesi *et al*., 2022), thus it is possible that this group performs the biosynthesis of ether lipids utilizing alternative biochemical strategies.

Recently, a radical SAM protein, a glycerol monoalkyl glycerol tetraether (GMGT) synthase (Gms), has been confirmed to be responsible for cross-linking the two hydrocarbon tails of isoprenoidal GDGT to produce isoprenoidal GMGTs (Garcia *et al*., 2024). The study by Garcia *et al*., 2024 also detected the presence of Gms homologs in bacterial genomes of the Acidobacteria, Planctomycetes, and diverse bacterial candidate phyla, and suggested that those homologs could be involved in the formation of the H covalent bond also in bacterial brGDGTs to generate brGMGTs. To this end and considering the detection of brGMGT-1a (*m/z* 1020) in our dataset, we also screened for the presence of homologs of the Gms protein in the Black Sea metagenomes. Most of the detected Gms homologs were affiliated with the phylum Desulfobacterota (84 / 262 MAGs) with maxima in the upper suboxic and deep euxinic waters, and with different groups of the Planctomycetota (90 / 684 MAGs) with higher abundance in the deeper suboxic (130-170 m) and euxinic waters (2,000 m) (Fig. S9, S10C Table S8, S15AB). The abundance of Gms homologs from Planctomycetota coincides with the distribution of brGMGT-1a (*m/z* 1020) (Fig. 5AB) with maxima at around 130 m, suggesting these Planctomycetota groups might be responsible for the synthesis of these lipids.

Surprisingly, hits of Mss, Ger, ElbD, and Agps were also detected in archaeal MAGs. Mss hits were only detected in the class Woesearchaeia (DPANN superphylum) with maximum abundance from 100 to 130 m depth (Fig. 7C, S19A), which also partially coincided with the distribution of the Ger and ElbD hits found in some DPANN Woesearchaeia MAGs (Fig. S9BC), suggesting that members of this DPANN group could potentially be synthesizing or modifying membrane lipids to make bacterial membrane spanning ether-based lipids in the suboxic zone. Although hypothetical, this would be exciting as it would be further supporting a potential synthesis of bacterial-like lipids by a DPANN archaeal group (Castelle *et al*., 2021; Villanueva *et al*., 2021), likely coming from a bacterial host as they do not have the biosynthetic genetic capacity to make the building blocks themselves (Castelle *et al*., 2018; Dombrowski *et al*., 2019, 2020).

#### Conclusions

By using a combination of metagenomics and targeted lipid analysis by high resolution accurate mass spectrometry in environmental samples, we have been able to better connect specific membrane lipids with their biological sources. We were able to identify previously unreported taxa in the Black Sea (i.e. Altiarchaea and Aenignmarchaea), as well as a high diversity of brGDGTs and OB-GDGTs, including GMGTs lipids.

In the euxinic waters, archaea of the Thermoplasmatota may contribute to the production of isoGDGT with rings while previously members of ANME-1 were expected to be the only source. This might have an effect on the use of isoGDGTs with 1 or 2 rings as biomarkers of anaerobic methane oxidation. Besides, we also suggest Planctomycetota, Cloacimonadota, Desulfobacterota, Chloroflexota, Actinobacteria, and Myxococcota as potential producers of brGDGTs and/or OB-GDGTs based on the detection of MSL and ether-forming coding genes in their MAGs. Members of these groups would make ideal candidates for cultivation attempts and physiological experiments to further investigate their membrane lipid composition. There are major cultivation challenges to overcome to isolate high pressure anaerobic microorganisms and directly study their lipids, but several successful attempts have been published in recent years (Van Vliet *et al*., 2020; Yadav *et al*., 2021). By applying reverse metagenomics and understanding the metabolic potential of these taxonomic groups, it is also possible to design novel cultivation strategies for groups of interest, like Thermoplasmatota or Bathyarchaeaia (Cross *et al*., 2019; Sun *et al*., 2020; Lewis *et al*., 2021).

Based on the detection of homologs of the archaeal *tes* and *grsA* genes, we also suggest bacteria of the Planctomycetota and Cloacimonadota could be synthesizing membrane spanning lipids potentially with cyclopentane rings. Confirming this hypothesis would be essential to elucidate how the ‘lipid divide’ occurred (Koga, 2011; Lombard *et al*., 2012; Villanueva *et al*., 2017, 2021; Sahonero-Canavesi *et al*., 2022), and if the physiological advantage of harboring membranes with mixed archaeal and bacterial features is to increase resilience and adaptability to environmental stress.

We also detected homolog of the bacterial MSL and ether-forming coding genes in Archaea, specifically in members of the DPANN Woesearchaea. Due to the restricted genomic potential this group possesses to produce their own lipids (Castelle *et al*., 2018; Dombrowski *et al*., 2019, 2020), this might suggest their ability to modify already formed membrane lipids taken from a bacterial host and the capability to make bacterial membrane spanning ether-lipids. Identifying the prospective hosts of Woesearchaea in euxinic waters and the isolations of host-symbiont pairs could lead to a better understanding of the physiological and evolutionary dynamics of these archaea.

#### Experimental procedures

##### Samples

Suspended particulate matter (SPM) from 15 depths across the water column (50– 2000 m) was collected at sampling station 2 (N42°53.8′, E30°40.7′, 2107 m depth) during the Phoxy cruise 64PE371 (BS2013) on 9–10 June 2013 as previously described (Villanueva *et al*., 2021). SPM was collected with McLane WTS-LV in situ pumps (McLane Laboratories Inc., Falmouth) on pre-combusted glass fiber filters with 142-mm diameter and 0.7-µm pore size.

##### Lipid analysis

Total lipids were extracted from freeze-dried glass fiber filters as described in Sollai *et al*., (2019). The Bligh and Dyer lipid extracts were analyzed for archaeal GDGTs intact polar lipids (IPLs), which are composed of the core lipid (CL) attached to one or two polar head groups, by Ultra High-Pressure Liquid Chromatography (UHPLC) system (an Agilent 1290 Infinity I) coupled to a Q Exactive Orbitrap high-resolution mass spectrometry (HRMS) system (Thermo Fisher Scientific, Waltham, MA). UHPLC–atmospheric pressure chemical ionization MS for archaeol and isoGDGTs, was done according to Hopmans *et al*., (2016) with some modifications as described in Sollai *et al*., (2019). SPM samples were re-extracted also following the Bligh and Dyer lipid extraction procedure but with extractions of the residue with a mixture of methanol, dichloromethane, and aqueous trichloroacetic acid solution (TCA) pH 3 (2:1:0.8, v:v) as described in Bale *et al*., (2021). These extracts were analyzed for brGDGTs and OB- GDGTs core lipids by using UHPLC-HRMS according to the reversed phase method of Wörmer *et al*., (2013) with the modifications by Bale *et al*., (2021).

A range of branched GDGTs (brGDGTs) were detected (Fig. S1, S4 for structures and nomenclature). Figure S4 shows the distribution of brGDGTs in SPM from 1000 m depth. The identification of all the brGDGT components was based on their accurate masses (Table S7) and by comparison with published fragmentation spectra (Liu, Summons, *et al*., 2012; Naafs *et al*., 2018, 20; Baxter *et al*., 2019). Further confirmation of the assignments, and in particular their elution order, came from re-examination of the brGDGT distribution in extracts from Lake Challa, Kenya (data not shown), under both normal phase analysis (Baxter *et al*., 2019) and reverse phase analysis (Mitrović *et al*., 2023). This allowed us to confirm the elution of brGMGT-1a (a “H-GDGT”, also known as brGDGT H1020) relative to brGDGT-1a. Indeed, the elution of brGMGT-1a during this study, under reverse phase analysis, was somewhat unexpected, forming a double peak (presumably two isomers) with maxima eluting around 0.4 and 0.7 minutes after that of brGDGT-1a (*mz* 1022). Under normal phase analysis (e.g.,Baxter *et al*., 2019) brGMGTs elute 20 minutes after their equivalent brGDGTs. Furthermore, during reverse phase analysis, isoprenoidal GMGTs elute a few minutes before their equivalent GDGTs (Mitrović *et al*., 2023). Several late eluting, overly branched (OB)-GDGTs (Liu, Summons, *et al*., 2012) were also detected (Fig. S4; Table S7), identified from their accurate masses and their elution pattern.

##### DNA extraction, 16S rRNA gene amplicon sequencing, and quantitative PCR

The DNA extraction from SPMs across the water column of the Black Sea was reported in Sollai *et al*., (2019). Microbial diversity determination by 16S rRNA gene amplicon sequencing was performed as described in (Villanueva *et al*., 2021). Quantitative PCR was performed using the same primers of the 16S rRNA gene amplicon sequencing as described in Villanueva *et al*., (2021)

##### Metagenomic sequencing, assembly and binning

The metagenomic sequencing, assembly, and binning is described and reported in Ding and von Meijenfeldt *et al*. (2024). In short, the 15 samples were individually assembled, sequencing reads were mapped back to the assembled scaffolds generating 15 x 15 mappings, and the scaffolds were binned per sample into 5,844 metagenome-assembled genomes (MAGs). MAG quality was assessed with CheckM v1.1.3105 (Parks *et al*., 2015) in the lineage-specific workflow, and the MAGs were taxonomically annotated with GTDB-Tk v2.1.0 (Chaumeil *et al*., 2022) using release 207 of the GTDB (Parks *et al*., 2022). The GTDB-Tk annotations were only used for the reconstruction of the archaeal phylogeny (see below) and in Fig. 2A and Table S4. Taxonomy was also assigned to scaffolds and MAGs with Contig Annotation Tool (CAT) and Bin Annotation Tool (BAT) (von Meijenfeldt *et al*., 2019), respectively, from the CAT pack software suite v5.2.3, using a reference database based on the set of non-redundant proteins in GTDB release 207. The CAT and BAT classifications were used in all other analyses. Prodigal v2.6.3 in metagenomic mode (Hyatt *et al*., 2012) and DIAMOND v2.0.6109 (Buchfink *et al*., 2021) were used for protein prediction and alignment, respectively. The --top parameter was set to 11 for CAT and to 6 for BAT. A MAG may get multiple taxonomic annotations with the default parameter settings of BAT (-f 0.3) in which case we took the majority classification (-f 0.5).

##### Archaeal phylogeny

A phylogeny was constructed representing the archaeal MAGs detected in the Black Sea and a set of reference genomes picked from across the archaeal tree of life with a high taxonomic sampling around the taxonomic groups that were detected in the Black Sea. Taxonomy of archaeal MAGs was assessed based on the GTDB-Tk annotations (see above). Similar MAGs in the Black Sea were identified with dRep v3.4.0 (Olm *et al*., 2017) sing the fastANI algorithm (--P_ani 0.9, --S_ani 0.99) and only including MAGs with a minimum completeness of 50% (--completeness 50). We downloaded representative genomes from GTDB, selecting one archaeal genome per order, and all representative genomes of clades that contain the archaeal MAGs from the Black Sea. The representative genome that was chosen per order was selected based on estimated CheckM completeness – 5 x contamination. Genes were identified with Prodigal v2.6.3 in single mode (Hyatt *et al*., 2010), and queried for the 27 Clusters of Orthologous Gene (COG) families that showed evidence of being primarily vertically transferred in (Moody *et al*., 2022), as described in (Gallego *et al*., 2024). 24 COG families were selected, and genes from these families that were present in single-copy in a genome were extracted and aligned with MAFT v7.505 (Katoh and Standley, 2013) using the L-INS-I algorithm (Katoh and Standley, 2013). Alignments were trimmed with trimAl v1.4.rev15 (Capella-Gutiérrez *et al*., 2009) in gappyout mode, and sequences that contained >60% gaps after trimming were removed. The aligned sequences were concatenated per genome, filling in gaps when a gene was absent from the alignment. The final concatenated alignment contained 208 representative MAGs from the Black Sea and 1,354 representative genomes from GTDB, and included 8,944 aligned amino acids.

We constructed a maximum-likelihood phylogenetic tree with IQ-TREE v2.1.2 (Nguyen *et al*., 2015) using 1,000 ultrafast bootstraps (Minh *et al*., 2013) and model selection (Kalyaanamoorthy *et al*., 2017) best-fit model (LG+F+R10) chosen based on the Bayesian Information Criterion (BIC). Visualization was done in Interactive Tree of Life (iTOL) (Letunic and Bork, 2024).

##### Search for homologs of targeted genes and their abundance profile

Homologs of the targeted genes were identified by blastp+ (v. 2.7.1) (Camacho and Madden, 2023) in all proteins predicted on the scaffolds by Prodigal with e-value < 1 x 10^-30^ and percentage of identity >= 30%. Query sequences of targeted protein coding genes were the following: Tes protein of *Methanococcus aeolicus* Nanakai-3 (accession number ABR56159.1), GrsA protein of *Sulfolobus acidocaldarius* (accession number WP_011278400.1), elbD protein of *Myxococcus xanthus* (accession number ABF88003.1), agps protein of *Myxococcus xanthus* (accession number ABF89845.1), Gms protein of *Thermococcus guaymasensis* DSM 11113 (accession number AJC70771.1).

Bubble plots for the visualization of hits in MAGs across the water column were done with Python 3 and Matplotlib in JupyterLab (https://jupyter.org/).

The script jgi_summarize_bam_contig_depths that comes with MetaBAT2 (Kang *et al*., 2019) was used to extract average depth (i.e. number of mapped reads per base pair) for each scaffold that contained hits from the BAM files, and depth was normalized per 1 x 10^8^ mapped reads in each sample. The CAT annotation of the scaffold, or the BAT annotation of the MAG if the scaffold was binned, was used for taxonomic classification of the hits.

The normalized average depth profiles of a scaffold that are based on all versus all mappings can be used to compare it to other scaffolds, but they cannot be summed to aggregate profiles from a certain taxon, as reads from one sample may map to similar strains in different samples, and thus be counted multiple times when summed. This leads to an overestimation of the abundance of taxa that are present in multiple samples. When aggregating the abundance of taxa, we thus summed normalized average depth only for those mappings from which the scaffold was assembled. For example, the summed normalized average depth of Halobacteriota with *grsA* homologs at 500 m depth only includes the scaffolds with hits assembled from that sample, and similarly for 1,000 m depth and 2,000 m depth. As no scaffolds of Halobacteriota with *grsA* hits were found in other samples, their summed normalized average depth is 0, even if the abundance profile of individual scaffolds (which is based on the all versus all mappings) may show presence in some of these samples.

## Data availability

The 16S rRNA gene amplicon reads (raw data) have been deposited in the NCBI Sequence Read Archive (SRA) under BioProject ID PRJNA423140, PRJNA649254–57. The Black Sea MAGs are deposited in Zenodo at https://doi.org/10.5281/zenodo.11453757. Phylogenetic tree file and iTOL annotation files are available in Zenodo at https://doi.org/10.5281/zenodo.12200552. Lipid data is available upon request by the reviewers and will be made publicly available.

## Acknowledgements

We thank Alejandro Abdala Asbun and Maartje Brouwer for their technical support. LV received funding from the Soehngen Institute for Anaerobic Microbiology (SIAM) through a Gravitation Grant (024.002.002) from the Dutch Ministry of Education, Culture, and Science (OCW). LV and DCB received funding from the ALW NWO Open Program ALWOP.256. FABvM received funding from a Spinoza award from NWO to Prof. Jaap Sinninghe Damsté.

## References

Bale, N.J., Ding, S., Hopmans, E.C., Arts, M.G.I., Villanueva, L., Boschman, C., et al. (2021) Lipidomics of Environmental Microbial Communities. I: Visualization of Component Distributions Using Untargeted Analysis of High-Resolution Mass Spectrometry Data. Front Microbiol 12: 659302.

Baxter, A.J., Hopmans, E.C., Russell, J.M., and Sinninghe Damsté, J.S. (2019) Bacterial GMGTs in East African lake sediments: Their potential as palaeotemperature indicators. Geochimica et Cosmochimica Acta 259: 155–169.

Baxter, A.J., Van Bree, L.G.J., Peterse, F., Hopmans, E.C., Villanueva, L., Verschuren, D., and Sinninghe Damsté, J.S. (2021) Seasonal and multi-annual variation in the abundance of isoprenoid GDGT membrane lipids and their producers in the water column of a meromictic equatorial crater lake (Lake Chala, East Africa). Quaternary Science Reviews 273: 107263.

Becker, K.W., Lipp, J.S., Versteegh, G.J.M., Wörmer, L., and Hinrichs, K.-U. (2015) Rapid and simultaneous analysis of three molecular sea surface temperature proxies and application to sediments from the Sea of Marmara. Organic Geochemistry 85: 42–53.

Blumenberg, M., Seifert, R., Reitner, J., Pape, T., and Michaelis, W. (2004) Membrane lipid patterns typify distinct anaerobic methanotrophic consortia. Proc Natl Acad Sci USA 101: 11111–11116.

Brocks, J.J. and Pearson, A. (2005) Building the Biomarker Tree of Life. Reviews in Mineralogy and Geochemistry 59: 233–258.

Buchfink, B., Reuter, K., and Drost, H.-G. (2021) Sensitive protein alignments at tree-of-life scale using DIAMOND. Nat Methods 18: 366–368.

Buckles, L.K., Villanueva, L., Weijers, J.W.H., Verschuren, D., and Damsté, J.S.S. (2013) Linking isoprenoidal GDGT membrane lipid distributions with gene abundances of ammoniaLoxidizing *T haumarchaeota* and uncultured crenarchaeotal groups in the water column of a tropical lake (L ake C halla, E ast A frica). Environmental Microbiology 15: 2445–2462.

Cabello-Yeves, P.J., Callieri, C., Picazo, A., Mehrshad, M., Haro-Moreno, J.M., Roda-Garcia, J.J., et al. (2021) The microbiome of the Black Sea water column analyzed by shotgun and genome centric metagenomics. Environmental Microbiome 16: 5.

Camacho, C. and Madden, T. (2023) BLAST+ Release Notes. In BLAST® Help [Internet]. National Center for Biotechnology Information (US).

Capella-Gutiérrez, S., Silla-Martínez, J.M., and Gabaldón, T. (2009) trimAl: a tool for automated alignment trimming in large-scale phylogenetic analyses. Bioinformatics 25: 1972–1973.

Castelle, C.J., Brown, C.T., Anantharaman, K., Probst, A.J., Huang, R.H., and Banfield, J.F. (2018) Biosynthetic capacity, metabolic variety and unusual biology in the CPR and DPANN radiations. Nat Rev Microbiol 16: 629–645.

Castelle, C.J., Méheust, R., Jaffe, A.L., Seitz, K., Gong, X., Baker, B.J., and Banfield, J.F. (2021) Protein Family Content Uncovers Lineage Relationships and Bacterial Pathway Maintenance Mechanisms in DPANN Archaea. Front Microbiol 12: 660052.

Chaumeil, P.-A., Mussig, A.J., Hugenholtz, P., and Parks, D.H. (2022) GTDB-Tk v2: memory friendly classification with the genome taxonomy database. Bioinformatics 38: 5315–5316.

Connock, G.T., Owens, J.D., and Liu, X.-L. (2022) Biotic induction and microbial ecological dynamics of Oceanic Anoxic Event 2. Commun Earth Environ 3: 136.

Cross, K.L., Campbell, J.H., Balachandran, M., Campbell, A.G., Cooper, C.J., Griffen, A., et al. (2019) Targeted isolation and cultivation of uncultivated bacteria by reverse genomics. Nat Biotechnol 37: 1314–1321.

Damsté, J.S.S., Hopmans, E.C., Pancost, R.D., Schouten, S., and Geenevasen, J.A.J. (2000) Newly discovered non-isoprenoid glycerol dialkylglycerol tetraether lipids in sediments. Chem Commun 1683–1684.

De Jonge, C., Hopmans, E.C., Stadnitskaia, A., Rijpstra, W.I.C., Hofland, R., Tegelaar, E., and Sinninghe Damsté, J.S. (2013) Identification of novel penta- and hexamethylated branched glycerol dialkyl glycerol tetraethers in peat using HPLC–MS2, GC–MS and GC–SMB-MS. Organic Geochemistry 54: 78–82.

De Jonge, C., Kuramae, E.E., Radujković, D., Weedon, J.T., Janssens, I.A., and Peterse, F. (2021) The influence of soil chemistry on branched tetraether lipids in mid- and high latitude soils: Implications for brGDGT- based paleothermometry. Geochimica et Cosmochimica Acta 310: 95– 112.

De Jonge, C., Stadnitskaia, A., Hopmans, E.C., Cherkashov, G., Fedotov, A., Streletskaya, I.D., et al. (2015) Drastic changes in the distribution of branched tetraether lipids in suspended matter and sediments from the Yenisei River and Kara Sea (Siberia): Implications for the use of brGDGT-based proxies in coastal marine sediments. Geochimica et Cosmochimica Acta 165: 200–225.

Ding, S., Meijenfeldt, F.A.B. von, Bale, N.J., Damsté, J.S.S., and Villanueva, L. (2024) Coupled metalipidomics-metagenomics reveal structurally diverse sphingolipids produced by a wide variety of marine bacteria. 2024.01.25.577268.

Dombrowski, N., Lee, J.-H., Williams, T.A., Offre, P., and Spang, A. (2019) Genomic diversity, lifestyles and evolutionary origins of DPANN archaea. FEMS Microbiology Letters 366:.

Dombrowski, N., Williams, T.A., Sun, J., Woodcroft, B.J., Lee, J.-H., Minh, B.Q., et al. (2020) Undinarchaeota illuminate DPANN phylogeny and the impact of gene transfer on archaeal evolution. Nat Commun 11: 3939.

Elling, F.J., Kattein, L., Naafs, B.D.A., Lauretano, V., and Pearson, A. (2023) Heterotrophic origin and diverse sources of branched glycerol monoalkyl glycerol tetraethers (brGMGTs) in peats and lignites. Organic Geochemistry 178: 104558.

Gallego, R.P., Meijenfeldt, F.A.B. von, Bale, N.J., Damsté, J.S.S., and Villanueva, L. (2024) Emergence and evolution of heterocyte glycolipid biosynthesis enabled specialized nitrogen fixation in cyanobacteria. 2024.05.17.594646.

Garcia, A.A., Chadwick, G.L., Liu, X.-L., and Welander, P.V. (2024) Identification of two archaeal GDGT lipid-modifying proteins reveals diverse microbes capable of GMGT biosynthesis and modification. Proc Natl Acad Sci U S A 121: e2318761121.

Günther, F., Thiele, A., Gleixner, G., Xu, B., Yao, T., and Schouten, S. (2014) Distribution of bacterial and archaeal ether lipids in soils and surface sediments of Tibetan lakes: Implications for GDGT-based proxies in saline high mountain lakes. Organic Geochemistry 67: 19–30.

Halamka, T.A., McFarlin, J.M., Younkin, A.D., Depoy, J., Dildar, N., and Kopf, S.H. (2021) ▪ Oxygen limitation can trigger the production of branched GDGTs in culture. Geochemical Perspectives Letters.

Hamersley, M.R., Lavik, G., Woebken, D., Rattray, J.E., Lam, P., Hopmans, E.C., et al. (2007) Anaerobic ammonium oxidation in the Peruvian oxygen minimum zone. Limnology & Oceanography 52: 923–933.

Hauptfeld, E., Pappas, N., van Iwaarden, S., Snoek, B.L., Aldas-Vargas, A., Dutilh, B.E., and von Meijenfeldt, F.A.B. (2024) Integrating taxonomic signals from MAGs and contigs improves read annotation and taxonomic profiling of metagenomes. Nat Commun 15: 3373.

Hopmans, E.C., Schouten, S., and Sinninghe Damsté, J.S. (2016) The effect of improved chromatography on GDGT-based palaeoproxies. Organic Geochemistry 93: 1–6.

Hyatt, D., Chen, G.-L., LoCascio, P.F., Land, M.L., Larimer, F.W., and Hauser, L.J. (2010) Prodigal: prokaryotic gene recognition and translation initiation site identification. BMC Bioinformatics 11: 119.

Hyatt, D., LoCascio, P.F., Hauser, L.J., and Uberbacher, E.C. (2012) Gene and translation initiation site prediction in metagenomic sequences. Bioinformatics 28: 2223–2230.

Iasakov, T.R., Kanapatskiy, T.A., Toshchakov, S.V., Korzhenkov, A.A., Ulyanova, M.O., and Pimenov, N.V. (2022) The Baltic Sea methane pockmark microbiome: The new insights into the patterns of relative abundance and ANME niche separation. Marine Environmental Research 173: 105533.

Jackson, D.R., Cassilly, C.D., Plichta, D.R., Vlamakis, H., Liu, H., Melville, S.B., et al. (2021) Plasmalogen Biosynthesis by Anaerobic Bacteria: Identification of a Two-Gene Operon Responsible for Plasmalogen Production in *Clostridium perfringens*. ACS Chem Biol 16: 6–13.

Jain, S., Caforio, A., Fodran, P., Lolkema, J.S., Minnaard, A.J., and Driessen, A.J.M. (2014) Identification of CDP-Archaeol Synthase, a Missing Link of Ether Lipid Biosynthesis in Archaea. Chemistry & Biology 21: 1392–1401.

Kalyaanamoorthy, S., Minh, B.Q., Wong, T.K.F., von Haeseler, A., and Jermiin, L.S. (2017) ModelFinder: fast model selection for accurate phylogenetic estimates. Nat Methods 14: 587– 589.

Kang, D.D., Li, F., Kirton, E., Thomas, A., Egan, R., An, H., and Wang, Z. (2019) MetaBAT 2: an adaptive binning algorithm for robust and efficient genome reconstruction from metagenome assemblies. PeerJ 7: e7359.

Katoh, K. and Standley, D.M. (2013) MAFFT Multiple Sequence Alignment Software Version 7: Improvements in Performance and Usability. Molecular Biology and Evolution 30: 772–780.

Kirkels, F.M.S.A., Usman, M.O., and Peterse, F. (2022) Distinct sources of bacterial branched GMGTs in the Godavari River basin (India) and Bay of Bengal sediments. Organic Geochemistry 167: 104405.

Koga, Y. (2011) Early Evolution of Membrane Lipids: How did the Lipid Divide Occur? J Mol Evol 72: 274–282.

Koga, Y. and Morii, H. (2007) Biosynthesis of Ether-Type Polar Lipids in Archaea and Evolutionary Considerations. Microbiol Mol Biol Rev 71: 97–120.

Letunic, I. and Bork, P. (2024) Interactive Tree of Life (iTOL) v6: recent updates to the phylogenetic tree display and annotation tool. Nucleic Acids Research gkae268.

Lewis, W.H., Tahon, G., Geesink, P., Sousa, D.Z., and Ettema, T.J.G. (2021) Innovations to culturing the uncultured microbial majority. Nat Rev Microbiol 19: 225–240.

Liu, X.-L., Lipp, J.S., Schröder, J.M., Summons, R.E., and Hinrichs, K.-U. (2012) Isoprenoid glycerol dialkanol diethers: A series of novel archaeal lipids in marine sediments. Organic Geochemistry 43: 50–55.

Liu, X.-L., Summons, R.E., and Hinrichs, K.-U. (2012) Extending the known range of glycerol ether lipids in the environment: structural assignments based on tandem mass spectral fragmentation patterns. Rapid Communications in Mass Spectrometry 26: 2295–2302.

Liu, X.-L., Zhu, C., Wakeham, S.G., and Hinrichs, K.-U. (2014) In situ production of branched glycerol dialkyl glycerol tetraethers in anoxic marine water columns. Marine Chemistry 166: 1–8.

Lloyd, C.T., Iwig, D.F., Wang, B., Cossu, M., Metcalf, W.W., Boal, A.K., and Booker, S.J. (2022) Discovery, structure and mechanism of a tetraether lipid synthase. Nature 609: 197–203.

Lombard, J., López-García, P., and Moreira, D. (2012) The early evolution of lipid membranes and the three domains of life. Nat Rev Microbiol 10: 507–515.

Lorenzen, W., Ahrendt, T., Bozhüyük, K.A.J., and Bode, H.B. (2014) A multifunctional enzyme is involved in bacterial ether lipid biosynthesis. Nat Chem Biol 10: 425–427.

Meador, T.B., Zhu, C., Elling, F.J., Könneke, M., and Hinrichs, K.-U. (2014) Identification of isoprenoid glycosidic glycerol dibiphytanol diethers and indications for their biosynthetic origin. Organic Geochemistry 69: 70–75.

von Meijenfeldt, F.A.B., Arkhipova, K., Cambuy, D.D., Coutinho, F.H., and Dutilh, B.E. (2019) Robust taxonomic classification of uncharted microbial sequences and bins with CAT and BAT. Genome Biology 20: 217.

Minh, B.Q., Nguyen, M.A.T., and von Haeseler, A. (2013) Ultrafast Approximation for Phylogenetic Bootstrap. Molecular Biology and Evolution 30: 1188–1195.

Mitrović, D., Hopmans, E.C., Bale, N.J., Richter, N., Amaral-Zettler, L.A., Baxter, A.J., et al. (2023) Isoprenoidal GDGTs and GDDs associated with anoxic lacustrine environments. Organic Geochemistry 178: 104582.

Moody, E.R., Mahendrarajah, T.A., Dombrowski, N., Clark, J.W., Petitjean, C., Offre, P., et al. (2022) An estimate of the deepest branches of the tree of life from ancient vertically evolving genes. eLife 11: e66695.

Naafs, B.D.A., McCormick, D., Inglis, G.N., and Pancost, R.D. (2018) Archaeal and bacterial H-GDGTs are abundant in peat and their relative abundance is positively correlated with temperature. Geochimica et Cosmochimica Acta 227: 156–170.

Nguyen, L.-T., Schmidt, H.A., von Haeseler, A., and Minh, B.Q. (2015) IQ-TREE: A Fast and Effective Stochastic Algorithm for Estimating Maximum-Likelihood Phylogenies. Molecular Biology and Evolution 32: 268–274.

Niemann, H., Stadnitskaia, A., Wirth, S.B., Gilli, A., Anselmetti, F.S., Sinninghe Damsté, J.S., et al. (2012) Bacterial GDGTs in Holocene sediments and catchment soils of a high Alpine lake: application of the MBT/CBT-paleothermometer. Climate of the Past 8: 889–906.

Olm, M.R., Brown, C.T., Brooks, B., and Banfield, J.F. (2017) dRep: a tool for fast and accurate genomic comparisons that enables improved genome recovery from metagenomes through de-replication. ISME J 11: 2864–2868.

Overmann, J., Abt, B., and Sikorski, J. (2017) Present and Future of Culturing Bacteria. Annu Rev Microbiol 71: 711–730.

Pancost, R.D. and Sinninghe Damsté, J.S. (2003) Carbon isotopic compositions of prokaryotic lipids as tracers of carbon cycling in diverse settings. Chemical Geology 195: 29–58.

Parks, D.H., Chuvochina, M., Rinke, C., Mussig, A.J., Chaumeil, P.-A., and Hugenholtz, P. (2022) GTDB: an ongoing census of bacterial and archaeal diversity through a phylogenetically consistent, rank normalized and complete genome-based taxonomy. Nucleic Acids Research 50: D785–D794.

Parks, D.H., Imelfort, M., Skennerton, C.T., Hugenholtz, P., and Tyson, G.W. (2015) CheckM: assessing the quality of microbial genomes recovered from isolates, single cells, and metagenomes. Genome Res 25: 1043–1055.

Pavlovska, M., Prekrasna, I., Dykyi, E., Zotov, A., Dzhulai, A., Frolova, A., et al. (2021) Niche partitioning of bacterial communities along the stratified water column in the Black Sea. MicrobiologyOpen 10: e1195.

Pearson, A., Flood Page, S.R., Jorgenson, T.L., Fischer, W.W., and Higgins, M.B. (2007) Novel hopanoid cyclases from the environment. Environmental Microbiology 9: 2175–2188.

Pearson, A. and Ingalls, A.E. (2013) Assessing the Use of Archaeal Lipids as Marine Environmental Proxies. Annual Review of Earth and Planetary Sciences 41: 359–384.

Rattray, J.E., Van De Vossenberg, J., Hopmans, E.C., Kartal, B., Van Niftrik, L., Rijpstra, W.I.C., et al. (2008) Ladderane lipid distribution in four genera of anammox bacteria. Arch Microbiol 190: 51– 66.

Rush, D. and Sinninghe Damsté, J.S. (2017) Lipids as paleomarkers to constrain the marine nitrogen cycle. Environmental Microbiology 19: 2119–2132.

Sahonero-Canavesi, D.X., Siliakus, M.F., Abdala Asbun, A., Koenen, M., Von Meijenfeldt, F.A.B., Boeren, S., et al. (2022) Disentangling the lipid divide: Identification of key enzymes for the biosynthesis of membrane-spanning and ether lipids in Bacteria. Sci Adv 8: eabq8652.

Schouten, S., Hopmans, E.C., Pancost, R.D., and Damsté, J.S.S. (2000) Widespread occurrence of structurally diverse tetraether membrane lipids: Evidence for the ubiquitous presence of low-temperature relatives of hyperthermophiles. Proceedings of the National Academy of Sciences 97: 14421–14426.

Schouten, S., Hopmans, E.C., and Sinninghe Damsté, J.S. (2013) The organic geochemistry of glycerol dialkyl glycerol tetraether lipids: A review. Organic Geochemistry 54: 19–61.

Sinninghe Damsté, J.S., Rijpstra, W.I.C., Foesel, B.U., Huber, K.J., Overmann, J., Nakagawa, S., et al. (2018) An overview of the occurrence of ether- and ester-linked iso-diabolic acid membrane lipids in microbial cultures of the Acidobacteria: Implications for brGDGT paleoproxies for temperature and pH. Organic Geochemistry 124: 63–76.

Sinninghe Damsté, J.S., Rijpstra, W.I.C., Hopmans, E.C., Foesel, B.U., Wüst, P.K., Overmann, J., et al. (2014) Ether- and Ester-Bound *iso* -Diabolic Acid and Other Lipids in Members of Acidobacteria Subdivision 4. Appl Environ Microbiol 80: 5207–5218.

Sinninghe Damsté, J.S., Rijpstra, W.I.C., Hopmans, E.C., Weijers, J.W.H., Foesel, B.U., Overmann, J., and Dedysh, S.N. (2011) 13,16-Dimethyl Octacosanedioic Acid (*iso* -Diabolic Acid), a Common Membrane-Spanning Lipid of Acidobacteria Subdivisions 1 and 3. Appl Environ Microbiol 77: 4147–4154.

Sohlenkamp, C. and Geiger, O. (2016) Bacterial membrane lipids: diversity in structures and pathways. FEMS Microbiology Reviews 40: 133–159.

Sollai, M., Villanueva, L., Hopmans, E.C., Reichart, G., and Sinninghe Damsté, J.S. (2019) A combined lipidomic and 16S RRNA gene amplicon sequencing approach reveals archaeal sources of intact polar lipids in the stratified Black Sea water column. Geobiology 17: 91–109.

Sun, Y., Liu, Y., Pan, J., Wang, F., and Li, M. (2020) Perspectives on Cultivation Strategies of Archaea. Microb Ecol 79: 770–784.

Suominen, S., Dombrowski, N., Sinninghe Damsté, J.S., and Villanueva, L. (2021) A diverse uncultivated microbial community is responsible for organic matter degradation in the Black Sea sulphidic zone. Environmental Microbiology 23: 2709–2728.

Tremblay, J., Singh, K., Fern, A., Kirton, E.S., He, S., Woyke, T., et al. (2015) Primer and platform effects on 16S rRNA tag sequencing. Front Microbiol 6:.

Tyler, J.J., Nederbragt, A.J., Jones, V.J., and Thurow, J.W. (2010) Assessing past temperature and soil pH estimates from bacterial tetraether membrane lipids: Evidence from the recent lake sediments of Lochnagar, Scotland. Journal of Geophysical Research: Biogeosciences 115:.

Tyson, G.W. and Banfield, J.F. (2005) Cultivating the uncultivated: a community genomics perspective. Trends in Microbiology 13: 411–415.

Van Vliet, D.M., Lin, Y., Bale, N.J., Koenen, M., Villanueva, L., Stams, A.J.M., and Sánchez-Andrea, I. (2020) Pontiella desulfatans gen. nov., sp. nov., and Pontiella sulfatireligans sp. nov., Two Marine Anaerobes of the Pontiellaceae fam. nov. Producing Sulfated Glycosaminoglycan-like Exopolymers. Microorganisms 8: 920.

Villanueva, L., Bastiaan Von Meijenfeldt, F.A., Westbye, A.B., Yadav, S., Hopmans, E.C., Dutilh, B.E., and Sinninghe Damsté, J.S. (2021) Bridging the membrane lipid divide: bacteria of the FCB group superphylum have the potential to synthesize archaeal ether lipids. The ISME Journal 15: 168–182.

Villanueva, L., Damsté, J.S.S., and Schouten, S. (2014) A re-evaluation of the archaeal membrane lipid biosynthetic pathway. Nat Rev Microbiol 12: 438–448.

Villanueva, L., Schouten, S., and Damsté, J.S.S. (2017) Phylogenomic analysis of lipid biosynthetic genes of Archaea shed light on the ‘lipid divide.’ Environmental Microbiology 19: 54–69.

van Vliet, D.M., von Meijenfeldt, F.A.B., Dutilh, B.E., Villanueva, L., Sinninghe Damsté, J.S., Stams, A.J.M., and Sánchez-Andrea, I. (2021) The bacterial sulfur cycle in expanding dysoxic and euxinic marine waters. Environmental Microbiology 23: 2834–2857.

Wakeham, S.G., Lewis, C.M., Hopmans, E.C., Schouten, S., and Sinninghe Damsté, J.S. (2003) Archaea mediate anaerobic oxidation of methane in deep euxinic waters of the Black Sea. Geochimica et Cosmochimica Acta 67: 1359–1374.

Wegener, G., Laso-Pérez, R., Orphan, V.J., and Boetius, A. (2022) Anaerobic Degradation of Alkanes by Marine Archaea. Annu Rev Microbiol 76: 553–577.

Weijers, J.W.H., Schouten, S., Hopmans, E.C., Geenevasen, J.A.J., David, O.R.P., Coleman, J.M., et al. (2006) Membrane lipids of mesophilic anaerobic bacteria thriving in peats have typical archaeal traits. Environmental Microbiology 8: 648–657.

Weijers, J.W.H., Schouten, S., Van Den Donker, J.C., Hopmans, E.C., and Sinninghe Damsté, J.S. (2007) Environmental controls on bacterial tetraether membrane lipid distribution in soils. Geochimica et Cosmochimica Acta 71: 703–713.

Welander, P.V., Coleman, M.L., Sessions, A.L., Summons, R.E., and Newman, D.K. (2010) Identification of a methylase required for 2-methylhopanoid production and implications for the interpretation of sedimentary hopanes. Proc Natl Acad Sci USA 107: 8537–8542.

Wörmer, L., Lipp, J.S., Schröder, J.M., and Hinrichs, K.-U. (2013) Application of two new LC–ESI–MS methods for improved detection of intact polar lipids (IPLs) in environmental samples. Organic Geochemistry 59: 10–21.

Xie, S., Liu, X.-L., Schubotz, F., Wakeham, S.G., and Hinrichs, K.-U. (2014) Distribution of glycerol ether lipids in the oxygen minimum zone of the Eastern Tropical North Pacific Ocean. Organic Geochemistry 71: 60–71.

Yadav, S., Koenen, M., Bale, N., Sinninghe Damsté, J.S., and Villanueva, L. (2021) The physiology and metabolic properties of a novel, low-abundance Psychrilyobacter species isolated from the anoxic Black Sea shed light on its ecological role. Environmental Microbiology Reports 13: 899–910.

Zeng, Z., Chen, H., Yang, H., Chen, Y., Yang, W., Feng, X., et al. (2022) Identification of a protein responsible for the synthesis of archaeal membrane-spanning GDGT lipids. Nat Commun 13: 1545.

Zeng, Z., Liu, X.-L., Farley, K.R., Wei, J.H., Metcalf, W.W., Summons, R.E., and Welander, P.V. (2019) GDGT cyclization proteins identify the dominant archaeal sources of tetraether lipids in the ocean. Proc Natl Acad Sci USA 116: 22505–22511.

Zeng, Z., Xiao, W., Zheng, F., Chen, Y., Zhu, Y., Tian, J., and Zhang, C. (2023) Enhanced production of highly methylated brGDGTs linked to anaerobic bacteria from sediments of the Mariana Trench. Front Mar Sci 10: 1233560.

