## Supplementary Figures for "Unraveling an unknown diversity of archaeal and bacterial tetraether membrane lipid producers in a euxinic marine system"

[1]*Royal Netherlands Institute for Sea Research. Department of Marine Microbiology and Biogeochemistry.*
[2]*Utrecht University. Faculty of Sciences. Department of Biology.*

**Figure S1.** Overview of the structures of isoprenoid glycerol dialkyl glycerol tetraethers (isoGDGTs), branched GDGTs (brGDGTs), overly branched GDGTs (OB-GDGTs) detected in this study and their mass-to-charge (*m/z*) ratio.

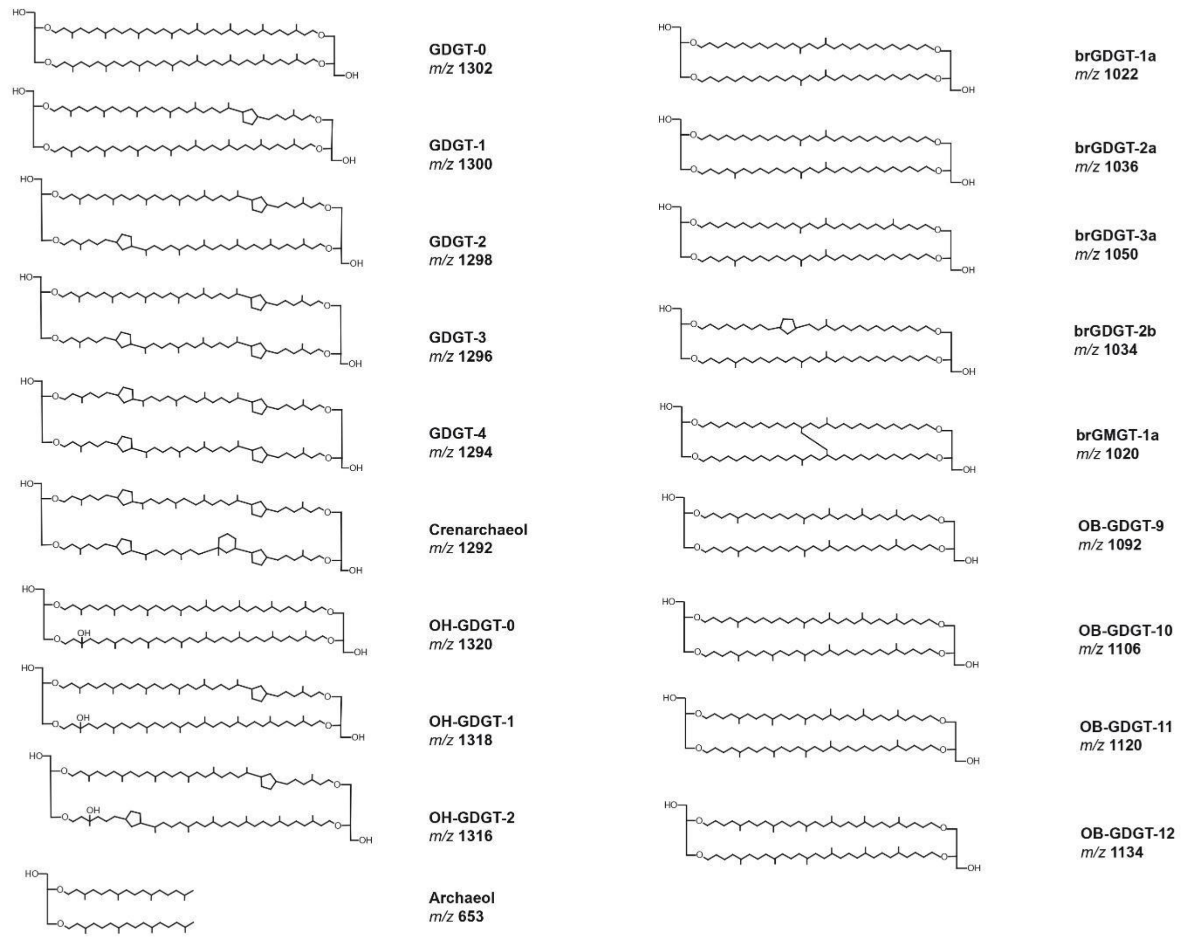

**Figure S2.** Overview of the physicochemical parameters of the Black Sea water column where the samples reported in this manuscript were collected. Data from Sollai *et al*., 2019.

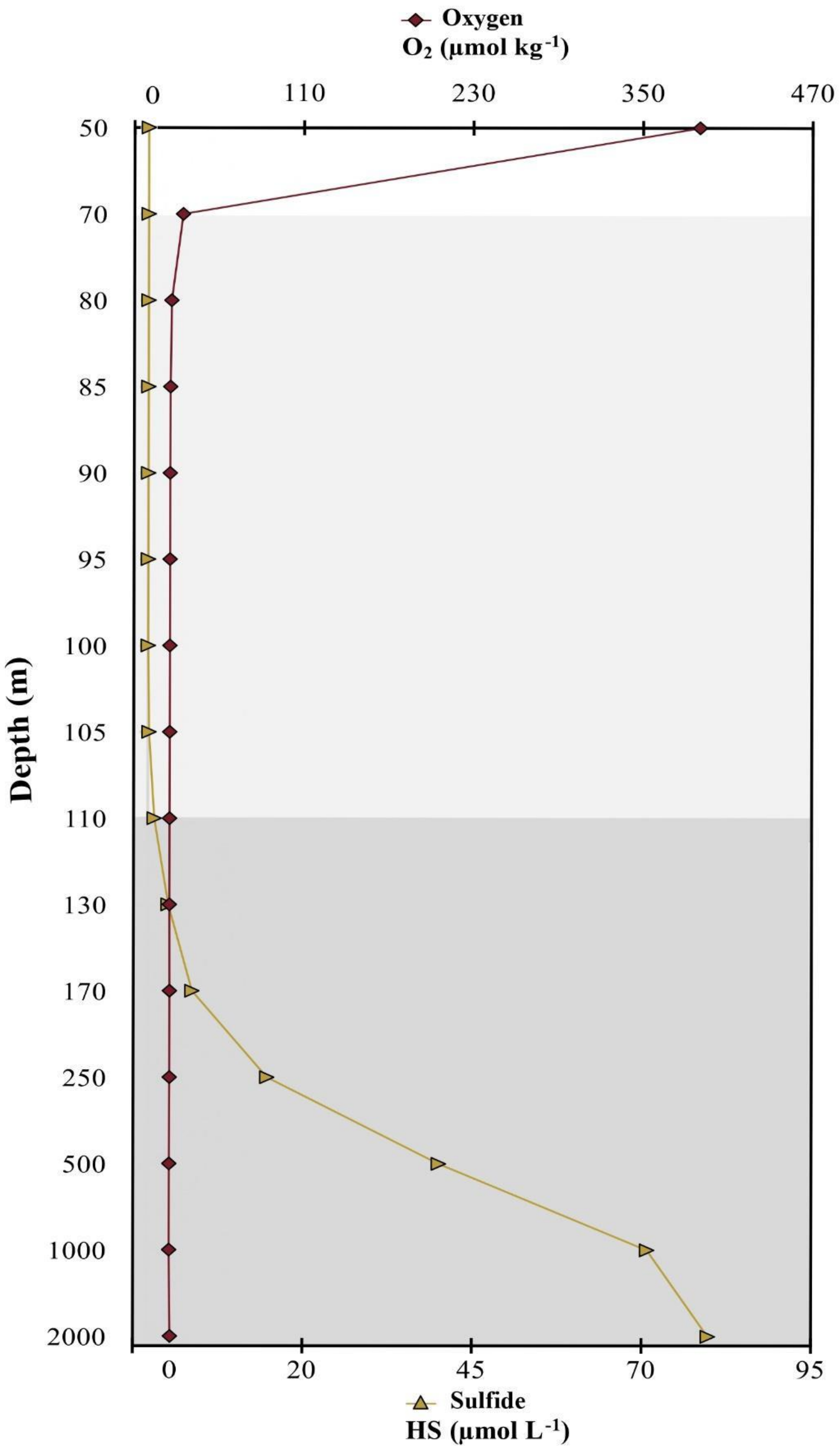

**Figure S3**. Overview of total archaeal and bacterial 16S rRNA gene reads per liter in the Black Sea water column suspended particulate matter (SPM) samples. Data from Sollai *et al*., 2019.

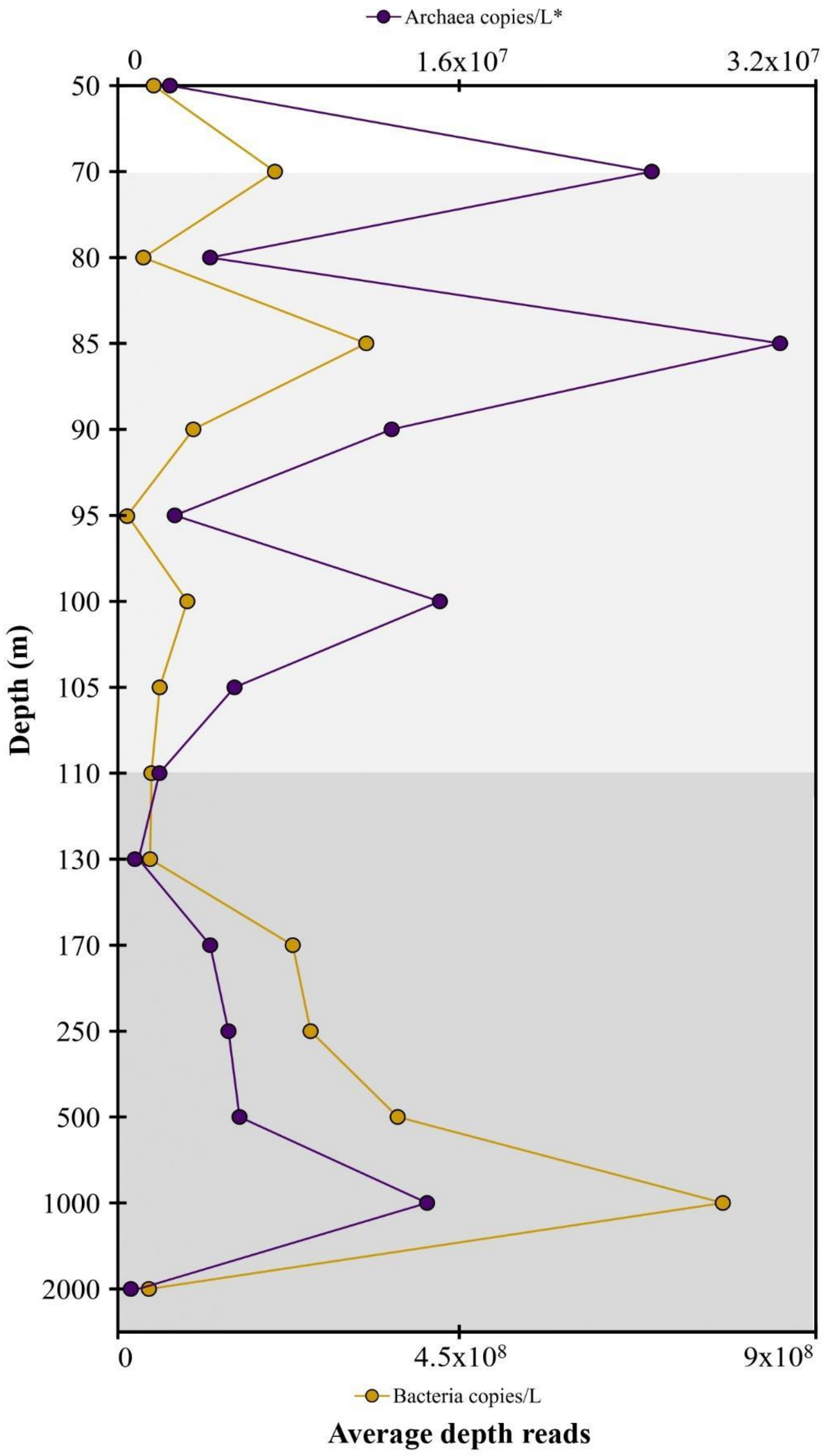

**Figure S4**. Extracted ion chromatograms (EICs) of branched (br) and overly branched (OB) GDGTs comprised between retention time 45.86-58.78 minutes, detected in the Black Sea 1,000 m suspended particulate matter sample and their accurate mass-to-charge (*m/z*) ratio (A-I, see main text for details).

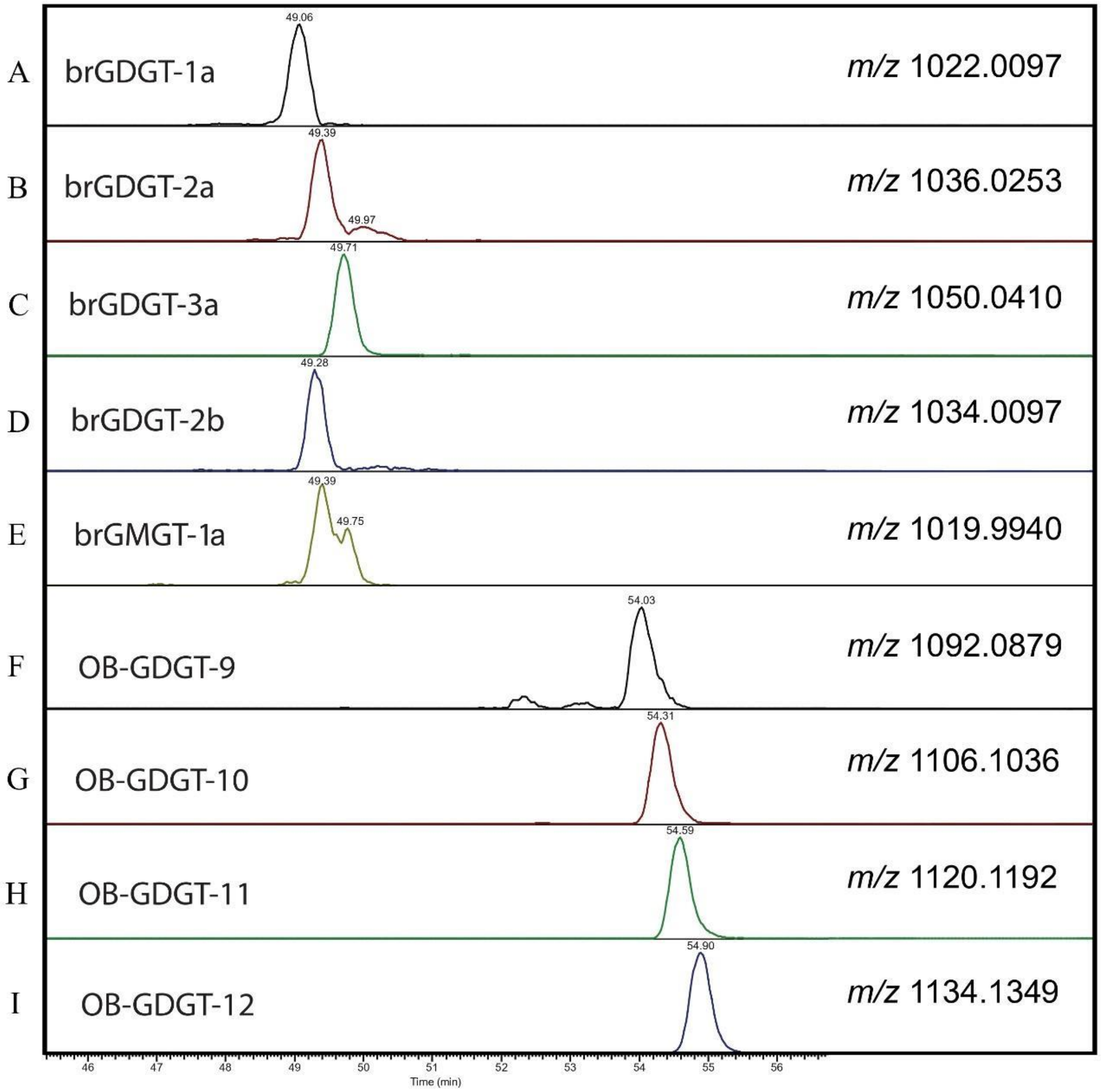

**Figure S5.** Sum of the average depth (i.e., number of mapped reads per base pair, per 1e+8 mapped reads) of (A) Tes protein of *Methanococcus aeolicus* Nanakai-3 (accession number ABR56159.1) homolog hits (protein blast e-value <= 1e-30, identity % >= 30%) and (B) GrsA protein of *Sulfolobus acidocaldarius* (accession number WP_011278400.1) homolog hits (protein blast e-value <= 1e-30, identity % >= 30%) detected in the different bacterial groups across the Black Sea SPM profile from 50 to 2,000 m depth. Data is compiled in Table S9AB, S10AB.

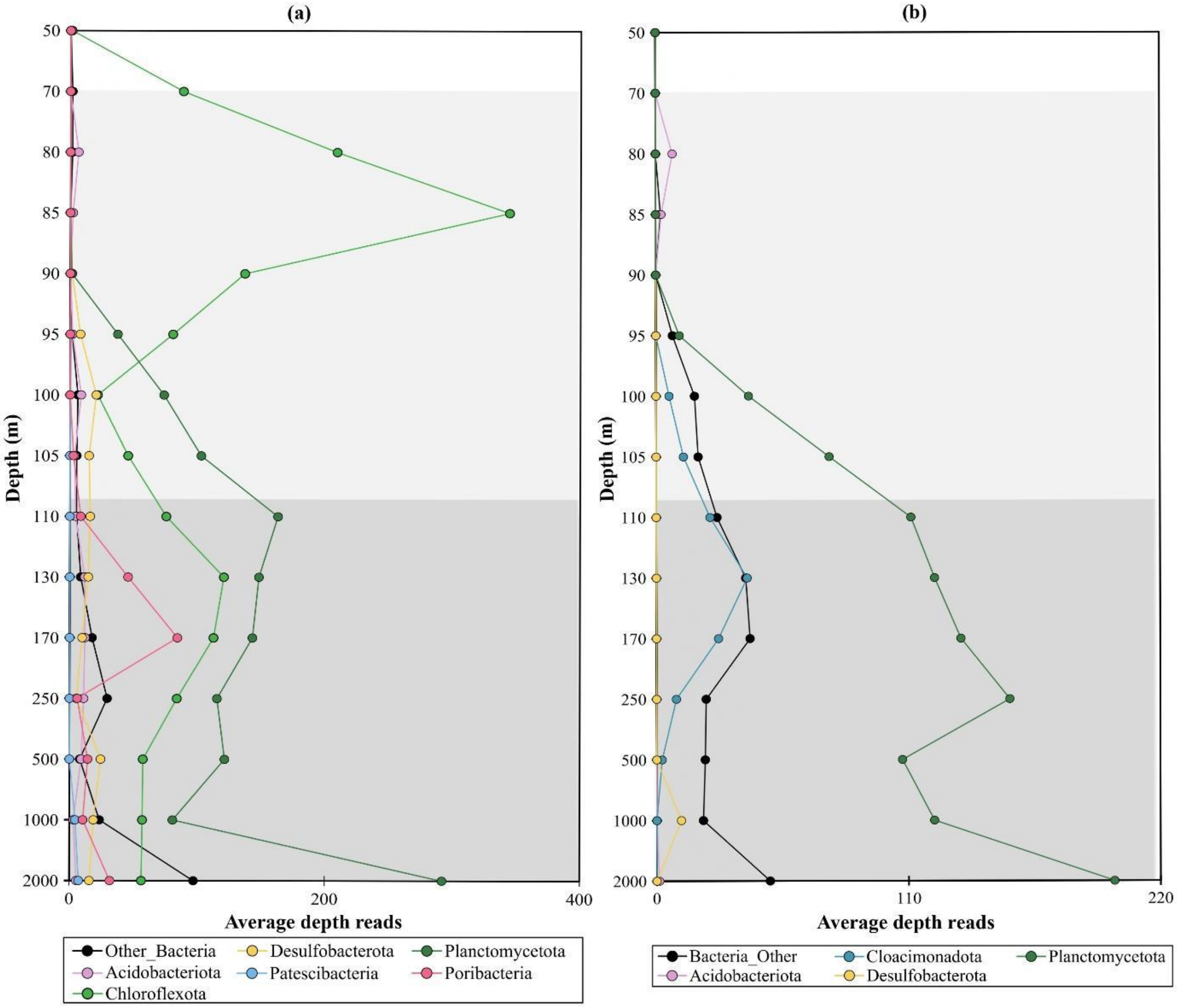

**Figure S6.** Sum of the average depth (i.e., number of mapped reads per base pair, per 1e+8 mapped reads) of ElbD protein of *Myxococcus xanthus* (accession number ABF88003.1) homolog hits (protein blast e-value <= 1e-30, identity % >= 30%) detected in the different bacterial groups across the Black Sea SPM profile from 50 to 2,000 m depth. Data is compiled in Table S13AB.

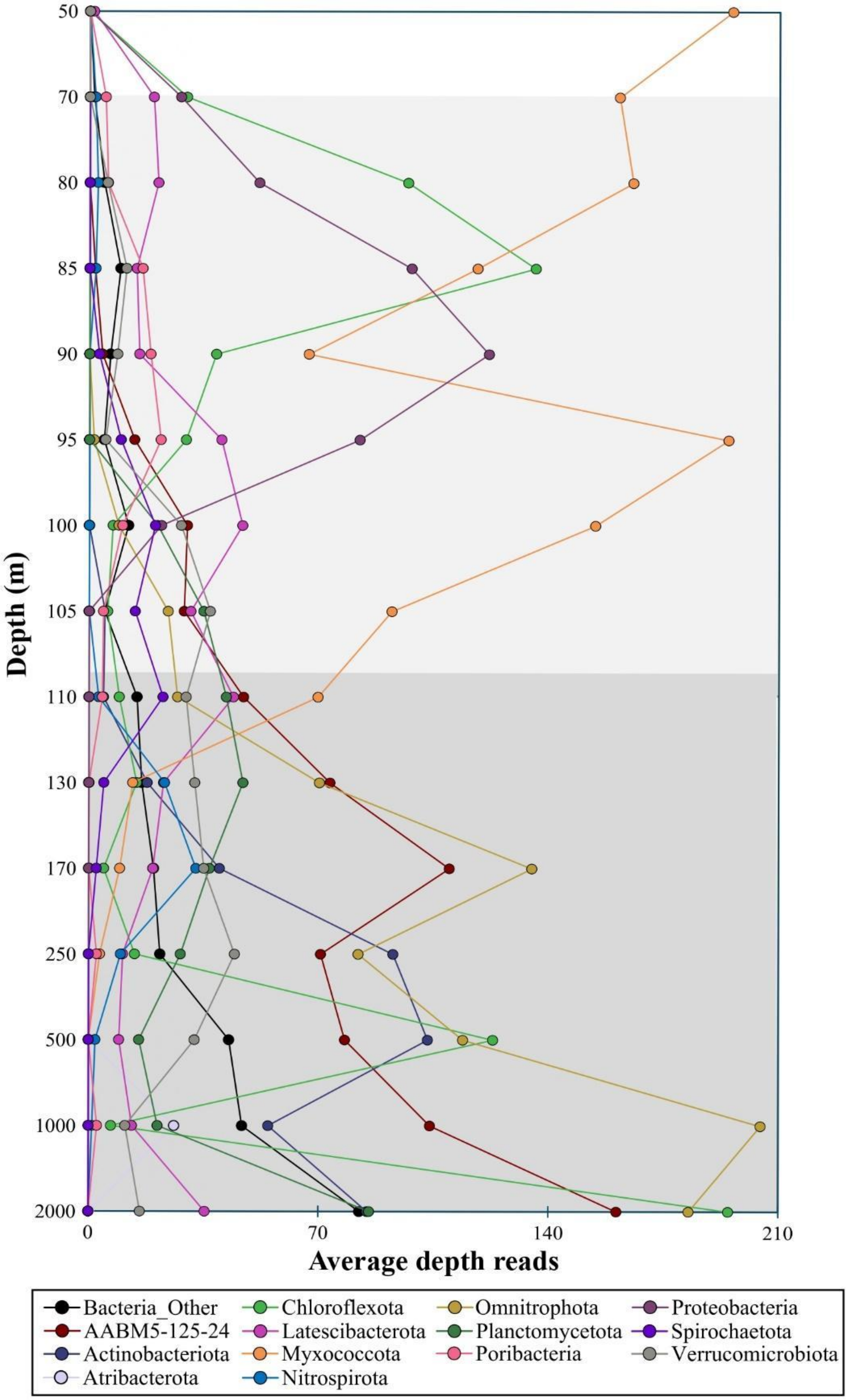

**Figure S7.** Sum of the average depth of the average depth (i.e., number of mapped reads per base pair, per 1e+8 mapped reads) of AgpsA protein of *Myxococcus xanthus* (accession number ABF89845.1) homolog hits (protein blast e-value <= 1e-30, identity % >= 30%) detected in the different bacterial groups across the Black Sea SPM profile from 50 to 2,000 m depth. Data is compiled in Table S14AB.

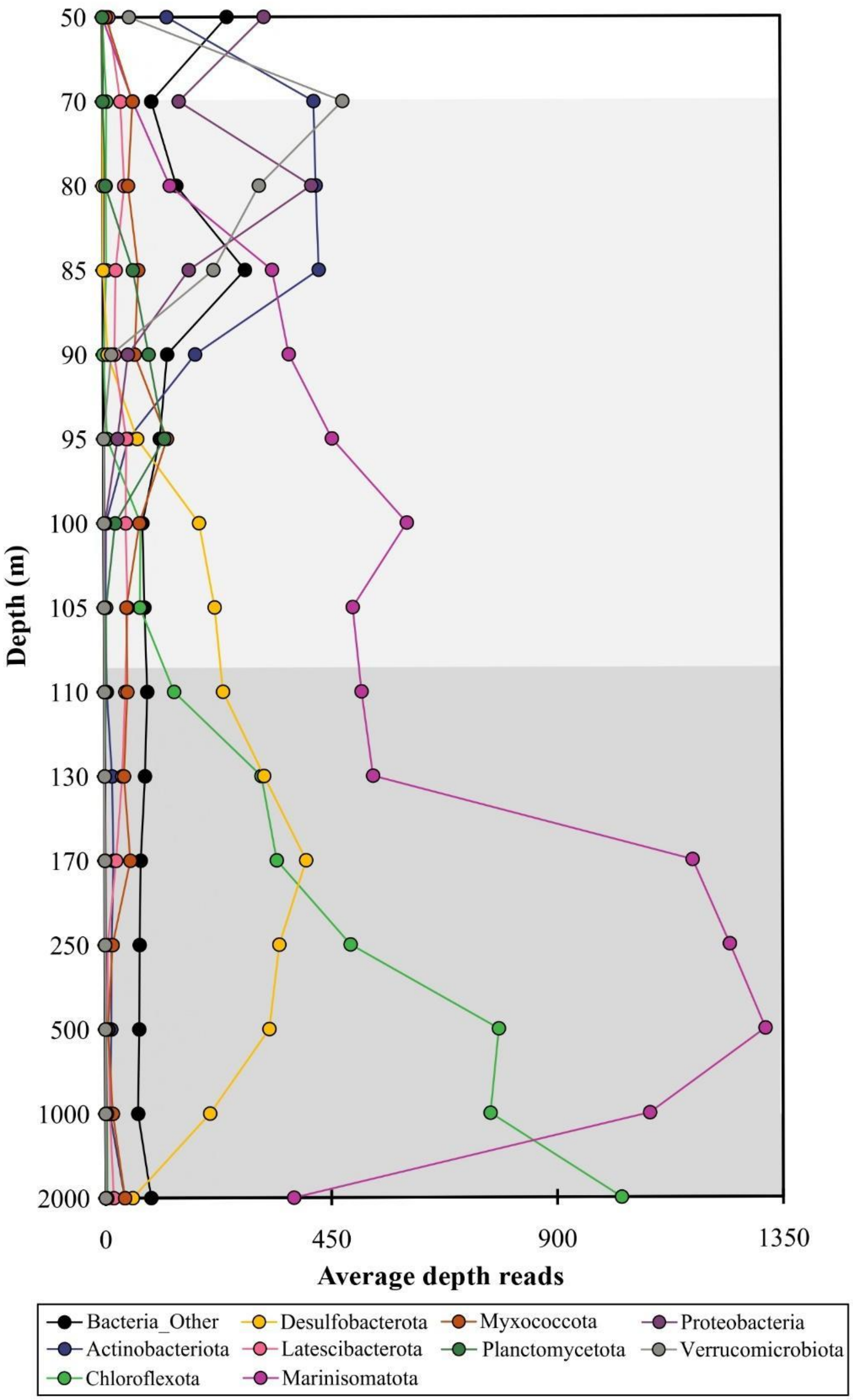

**Figure S8**. Sum of the average depth (i.e., number of mapped reads per base pair, per 1e+8 mapped reads) of Gms protein of *Thermococcus guaymasensis* DSM 11113 (accession number AJC70771.1) homolog hits (protein blast e-value <= 1e-30, identity % >= 30%) detected in the different microbial groups across the Black Sea SPM profile from 50 to 2,000 m depth. Data is compiled in Table S15AB.

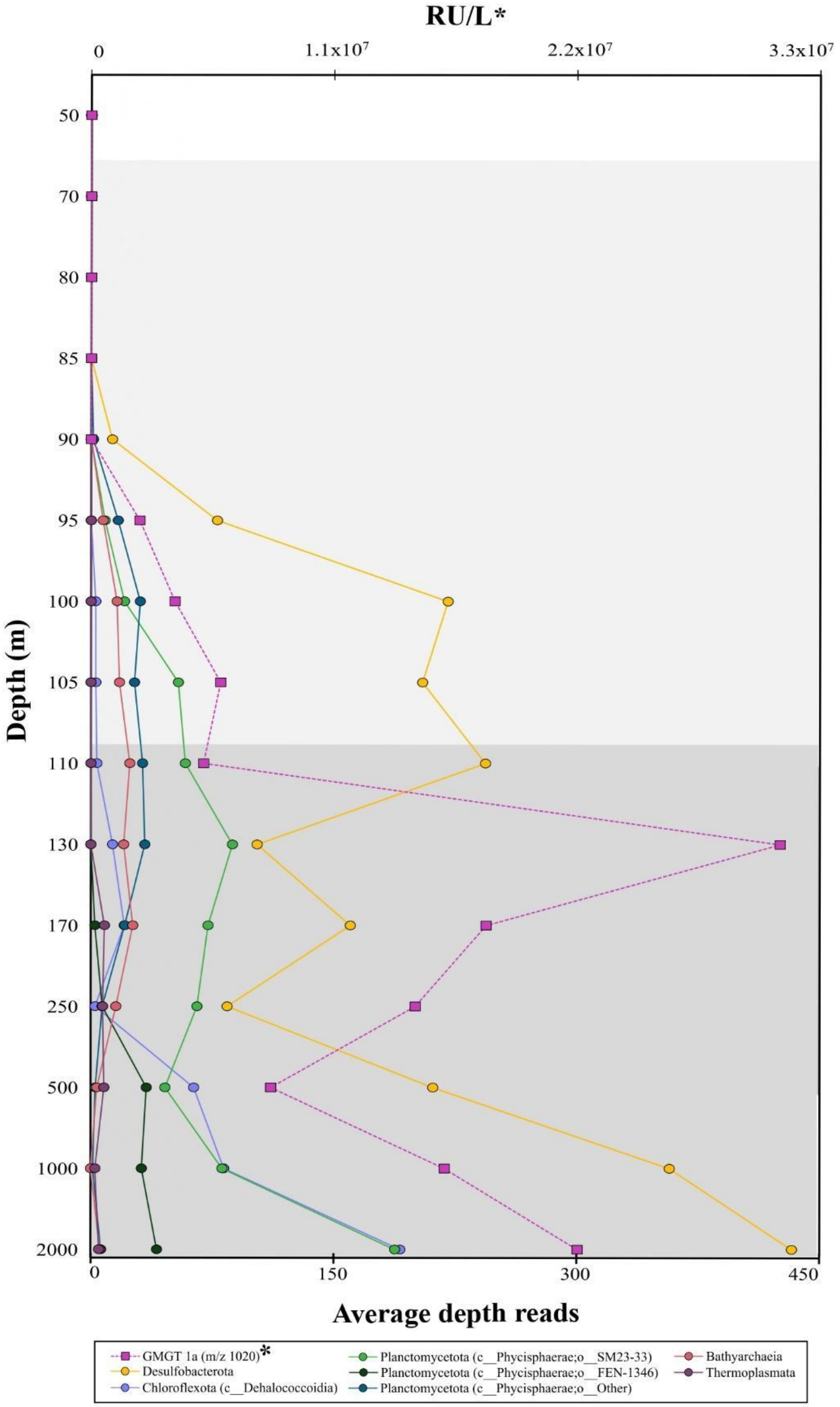


\

**Figure S9**. Distribution of the average depth (i.e., number of mapped reads per base pair, per 1e+8 maped reads) of Mss (a), Ger (b), ElbD (c) and Agps (d) homolog hits detected in the different archaeal groups across the Black Sea SPM profile from 50 to 2,000 m depth. Data is compiled in Tables S11AB, S12AB, S13AB, S14AB.

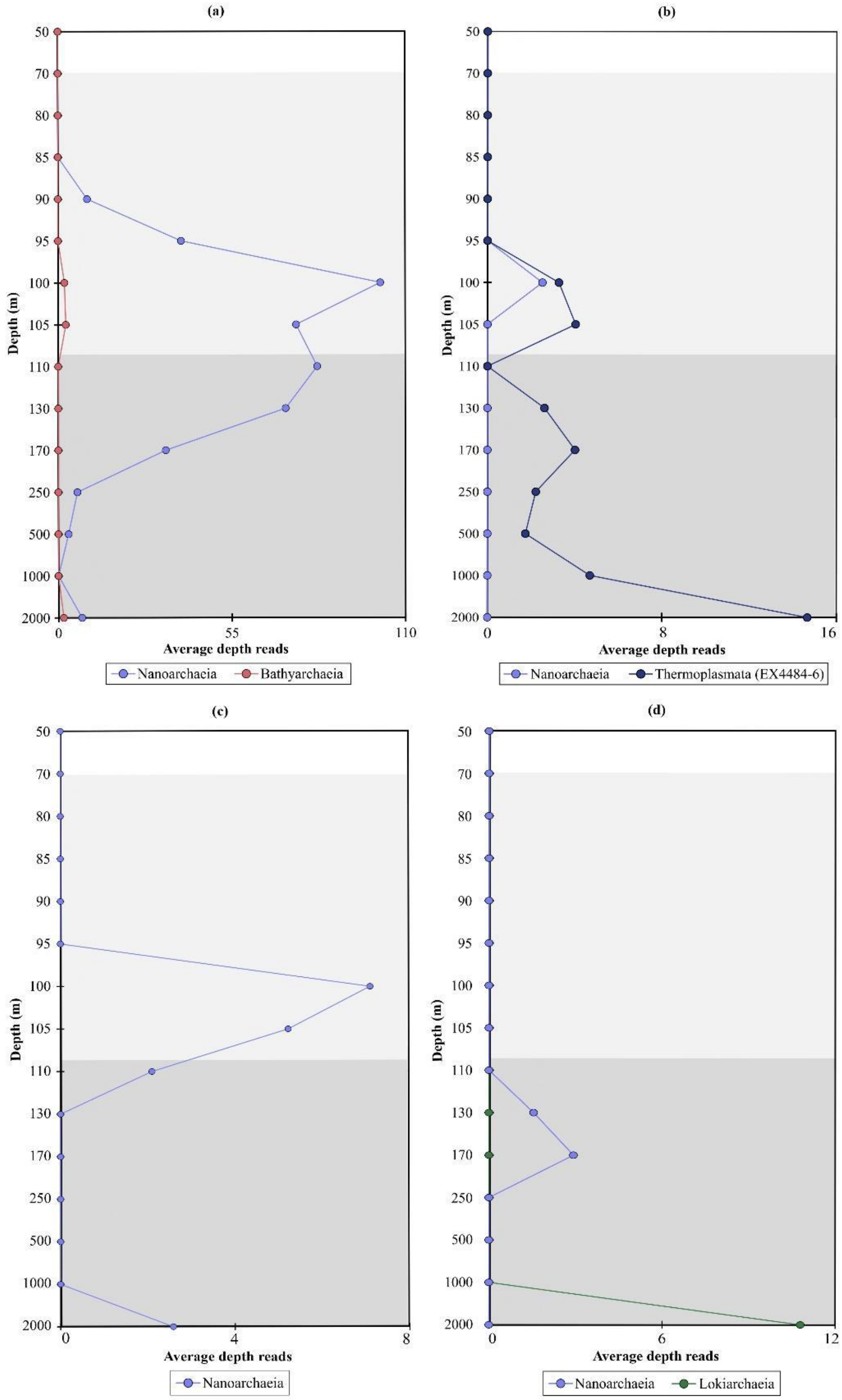

**Figure S10.** Number of metagenome-assembled genomes (MAGs) binned in the Black Sea aggregated by taxonomy (rank: phylum) and sample (depth). Circle size represents the number of MAGs by sample/taxonomy, color intensity describes fraction of MAGs containing genes: a)ElbD; b) Agps; c)Gms. Data is compiled in Tables S8.

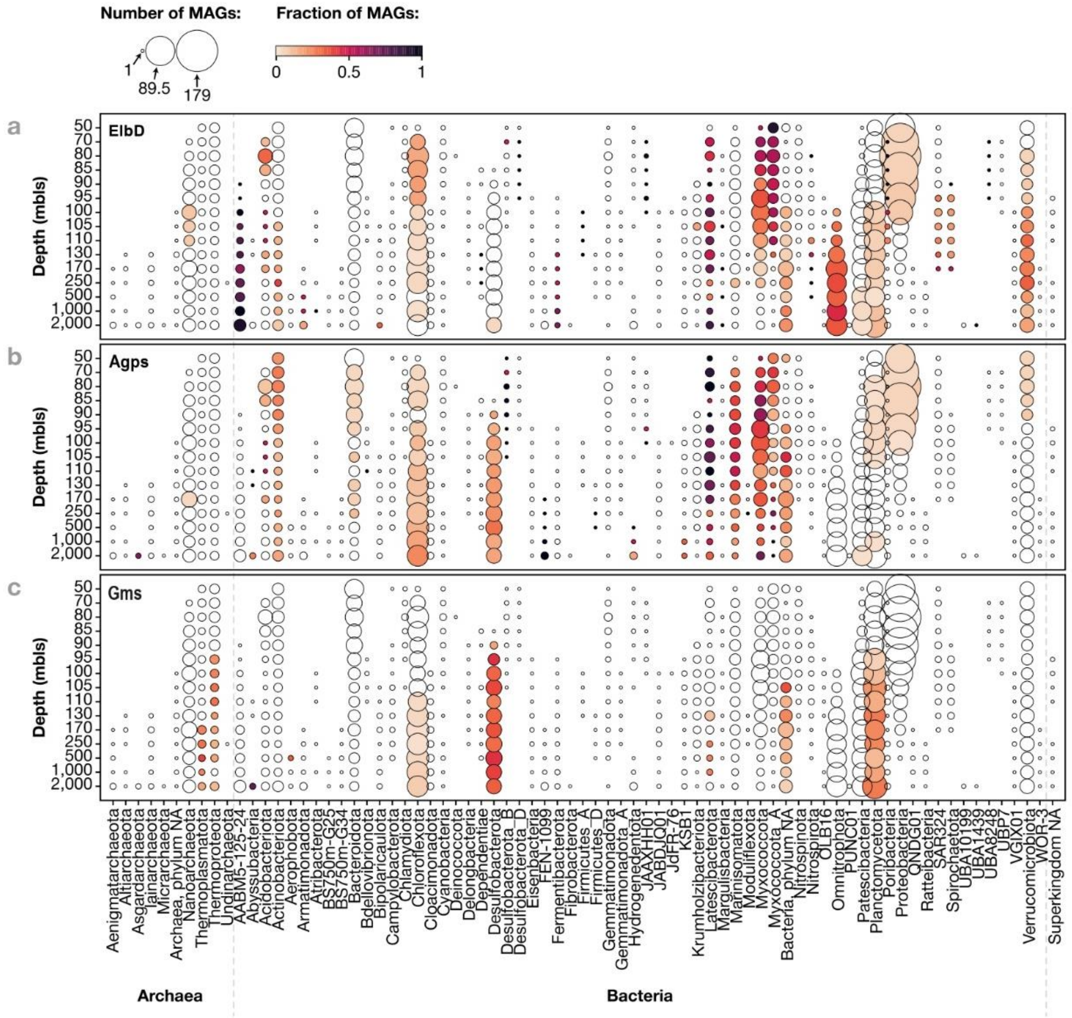
